# A Cajal body assembly factor regulates cell fate transitions in Arabidopsis

**DOI:** 10.64898/2026.08.19.745738

**Authors:** Jana Faturova, Vivek Kumar Raxwal, Marieke Dubois, Albert Cairo, Lucie Crhak-Khaitova, Sona Bukovcakova Valuchova, Matej Drs, Pavlina Mikulkova, Anna Vargova, Katerina Malinska, Lieven De Veylder, Karel Riha

## Abstract

Ribonucleoprotein (RNP) condensates are emerging as key regulators of cell fate transitions, yet their functions have been largely linked to mRNA storage and translational control. Here, we uncover a role for Cajal body (CB)-mediated pre-mRNA splicing in coordinating the transition from stem cell divisions to differentiation in plants. We identify THREE-DIVISION MUTANT 3 (TDM3) as a cell cycle-regulated factor required for post-mitotic CB assembly. Loss of TDM3 or the CB scaffold protein COILIN delays differentiation and prolongs formative cell divisions. Transcriptome analysis revealed that TDM3 and COILIN jointly regulate pre-mRNA splicing, including transcripts controlling cell cycle and fate transitions. These findings establish CB-mediated splicing as a mechanism linking cell cycle progression to cellular differentiation.

## Main

Cell fate transitions require extensive rewiring of gene expression. Although cellular reprogramming has been primarily attributed to transcriptional and chromatin-based mechanisms, post-transcriptional regulation is increasingly recognized as an additional important layer of control ^1–3^. Through coordinated control of RNA processing, localization, stability, and translation, post-transcriptional mechanisms enable rapid and reversible modulation of gene expression. This flexibility allows cells to fine-tune protein production and facilitate timely transitions between cellular states. Ribonucleoprotein (RNP) condensates provide a spatial framework for this coordination by compartmentalizing RNA-processing factors and dynamically regulating RNA metabolism. Accordingly, RNP condensates have been implicated in diverse reprogramming processes ^4–8^, yet their functions have largely been linked to mRNA storage and translational control.

Cellular differentiation is associated with extensive changes in splicing patterns and efficiency ^9–11^. Alternative splicing (AS) can regulate cell fate transitions by generating cell type-specific isoforms that either promote or restrain differentiation pathways ^12–14^. Beyond expanding proteomic diversity, AS can also fine-tune expression through mechanisms such as AS-coupled nonsense-mediated decay ^15^, which selectively degrades transcripts, and intron retention, which delays translation until a specific developmental stage ^16–19^. Shifts in splicing patterns may arise, at least in part, from increased transcriptional burden associated with cell state transitions, which places high demands on the splicing machinery ^16,20^. These observations suggest that developmental transitions may create transient bottlenecks in RNA processing, necessitating mechanisms that safeguard splicing capacity.

Cajal bodies (CBs) are nuclear ribonucleoprotein condensates that promote the biogenesis, maturation, and recycling of small nuclear ribonucleoproteins (snRNPs) and other factors required for splicing and RNA processing ^21–23^. By accelerating snRNP assembly ^24^, CBs become particularly important in highly active cells with elevated demands on RNA processing ^25^. However, whether CBs contribute directly to developmental reprogramming and cell fate transitions remains largely unknown.

Here, we identify a function of CBs in coordinating the timely transition of plant stem cells from proliferative divisions to differentiation. This developmental switch is often accompanied by a shift from mitotic cell cycles to endoreplication, which requires precise regulation of core cell-cycle components. We demonstrate that THREE-DIVISION MUTANT 3 (TDM3) is essential for a proper CB assembly during mitotic exit. TDM3 affects splicing of genes that control differentiation and endoreplication and its loss delays exit from mitotic proliferation. Our findings uncover a previously unrecognized role for CBs in developmental reprogramming and reveal how nuclear condensates couple pre-mRNA splicing to cell fate determination in plants

## Results

### Cell-cycle regulated expression of TDM3 in dividing cells

We have previously shown in *Arabidopsis thaliana* that the termination of meiosis and the transition to microspore differentiation are governed by M-bodies, specialized cytoplasmic RNP condensates consisting of a P-body core surrounded by a stress granule-like shell ^7,26^. This transition is facilitated by translational downregulation mediated by TDM1, a tetratricopeptide repeat protein that associates with M-bodies during meiosis II and sequesters the translation initiation factor eIFiso4G2 ^7^. TDM1 belongs to a plant specific family consisting of five members in Arabidopsis with similar AlphaFold predicted protein structure (Extended Data Fig. 1a)^27–29^. Among these, TDM3 (At3g51280) appears to be the most evolutionary ancient member of the family, with homologs present in liverworts and green algae, which lack the other TDM-like proteins (Fig. 1a and Extended Data Fig. 2).

**Fig. 1:**
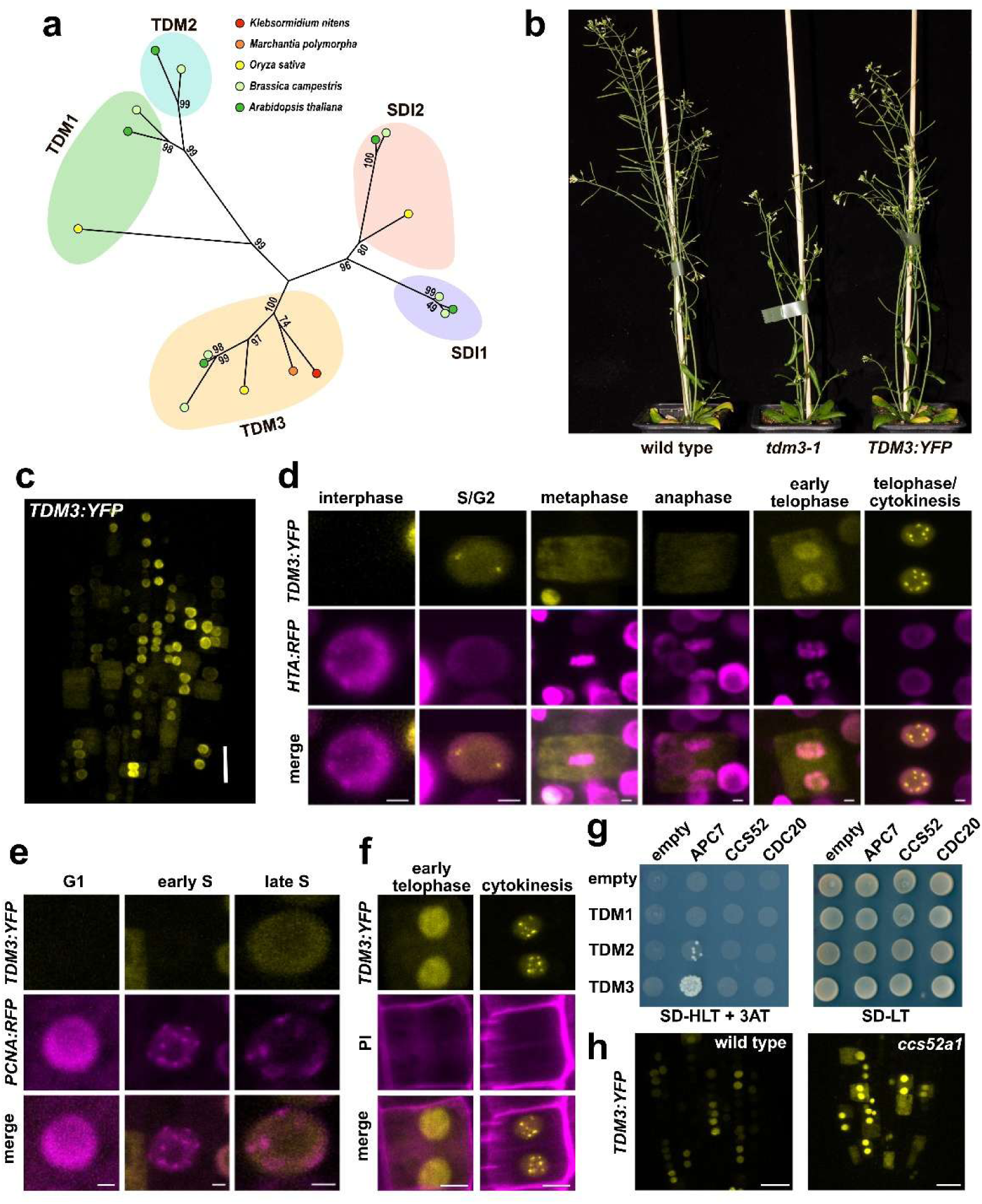
Cell-cycle dependent regulation of TDM3. a, Phylogenetic analysis of TDM/SDI homologs across representative plant species. Numbers at branch points indicate bootstrap support values from 1,000 replicates. **b,** Comparison of WT, *tdm3-1* and complemented *pTDM3::TDM3:YFP* plants. **c,** Representative maximum intensity projection (MIP) showing *pTDM3::TDM3:YFP* localization in the root meristem. **d,** Representative MIPs showing the localization of *pTDM3::TDM3:YFP* and the chromatin marker HTA:RFP in the course of cell cycle. **e,** Representative MIPs showing *pTDM3::TDM3:YFP* and the S-phase marker PCNA:RFP during G1-S phases. **f,** Representative MIPs showing *pTDM3::TDM3:YFP* and propidium iodide during telophase and cytokinesis. **g,** Interaction of TDM-like proteins with APC/C components analysed by yeast two-hybrid assay. SD-HLT supplemented with 2.5 mM 3-AT was used for interaction selection, and SD-LT served as growth control. **h,** MIPs of representative root apical meristems expressing *pTDM3::TDM3:YFP* in WT and *ccs52a1* mutants. The relative signal intensity of TDM3:YFP was 50±11 per nucleus in WT (n = 17) and 138±38 in *ccs52a1* plants (n = 15). Scale bars, 20 μm (**c,h**) and 2 μm (**d–f**).

Arabidopsis *tdm3-1* mutants carrying a T-DNA insertion in the first exon exhibit reduced stature and slower growth, indicating that, in contrast to TDM1, the function of TDM3 is not restricted to meiosis (Fig. 1b; Extended Data Fig. 1b). Analysis of *tdm3-1* plants complemented with the YFP-tagged *TDM3* gene driven by endogenous promoter (*TDM3:YFP*) revealed TDM3 expression in dividing cells across multiple tissues (Fig. 1b,c and Extended Data Fig. 3). In the root meristem, TDM3 displays a highly dynamic, cell-cycle dependent expression pattern. TDM3 accumulation initiates during S-phase, when the protein localizes to the nucleus and remains nuclear through G2 phase. Following nuclear envelope breakdown at the onset of M-phase, TMD3 distributes to cytoplasm. During telophase, TDM3:YFP relocalizes to the newly formed nuclei, where it transiently forms prominent nuclear speckles throughout cytokinesis, before rapidly disappearing upon completion of cell division (Fig. 1d-f, Videos 1 and 2). This expression pattern is consistent with the cell cycle regulated abundance of TDM3 mRNA, which peaks in M-phase in synchronized cell culture ^30^.

The sudden clearance of TDM3 at the end of M-phase suggests its degradation by the anaphase promoting complex/cyclosome (APC/C). Indeed, TMD3 contains a destruction box, a short motif recognized by APC/C co-activators, and directly interacts with the APC7 subunit (Fig. 1g, Extended Data Fig. 1 and Extended Data Fig.4a), which has been implicated in facilitating interactions between APC/C and its substrates ^31,32^.

Furthermore, TDM3:YFP expression in root cells increases upon treatment with proteasome inhibitor and in *ccs52a1-1* mutants deficient in the APC/C co-activator Cdh1 (Fig. 1h and Extended Data Fig. 4b)^33^. Collectively, these data show that TDM3 is a nuclear, cell-cycle regulated protein expressed from S to M-phases and degraded by APC/C after completion of cell division.

### TDM3 is required for proper post-mitotic Cajal body reassembly

To determine the identity of the nuclear TDM3 speckles, we performed colocalization studies in leaf mesophyll protoplasts transiently transfected with markers for different subnuclear compartments (Extended Data Fig. 5 and Fig. 2a). TDM3 signals co-localized with COILIN, the scaffold protein of Cajal bodies (CBs)^34,35^, and with the nucleolar protein fibrillarin (Fig. 2a). The co-localization of TDM3 with CBs was confirmed in plants co-expressing both *TDM3:YFP* and *COILIN:RFP* from their native promoters (Fig. 2b), whereas the nucleolar localization detected in protoplasts was not observed *in planta* and likely represents an artifact of overexpression. Localization of TDM3 to nuclear speckles was completely abolished in Arabidopsis *coilin-2* mutants (Fig. 2c; Extended Data Fig. 6). We further found that TDM3 and COILIN physically interact and that this interaction is specific to TDM3, as it was not detected with the closely related TDM1 protein (Fig. 2d,e). Together, these results establish TDM3 as a protein that interacts with COILIN and transiently associates with CBs at the end of mitosis.

**Fig. 2:**
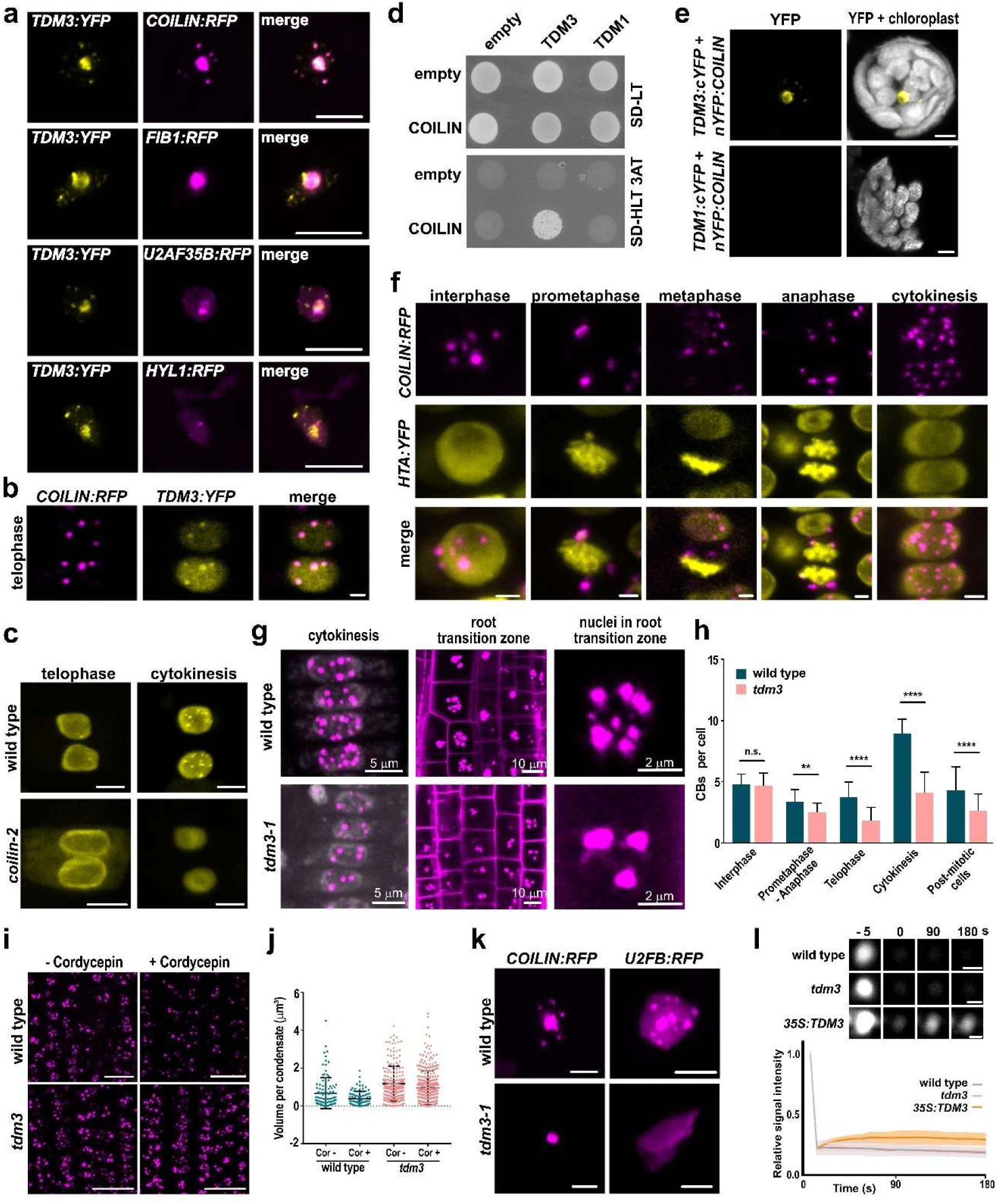
Role of TDM3 in mitotic Cajal body reassembly. a, Representative MIPs showing colocalization of TDM3 with nuclear compartment markers in mesophyll protoplasts. COILIN was used as a marker for Cajal bodies (CBs), FIBRILARIN1 (FIB1) for the nucleolus, U2AF35B for splicing speckles and HYL1 for dicing bodies. **b,** Co-localization of TDM3 and COLLIN during cytokinesis in roots expressing *pTDM3::TDM3:YFP* and *pCOILIN::COILIN:RFP*. **c,** Localization of *pTDM3::TDM3:YFP* during telophase and cytokinesis in WT and *coilin-2* in root nuclei. **d,** Yeast two-hybrid analysis interaction between TDM1 and TDM3 with COILIN. Interactions were assessed on SD-HLT medium, and SD-LT was used as growth control. **e,** BiFC assay in mesophyll protoplasts showing interaction between TDM3 and COILIN. **f,** Representative MIPs showing localization of CBs during mitosis in plants expressing *pCOILIN::COILIN:RFP* and the chromatin marker HTA:YFP. **g,** Localization of CBs during cytokinesis in root meristematic and postmitotic cells expressing *pCOILIN::COILIN:RFP*. **h,** Quantification of CBs number across cell-cycle stages in *tdm3-1* and WT. Data represent mean ± s.d.; two-sided Wilcoxon rank-sum tests with Holm correction were used for *\*\*P < 0.01, ****P < 0.0001*; n.s., not significant. n = biologically independent samples: WT, n = 8 roots and 20 cells for interphase, n = 8 roots and 18 cells for prometaphase–anaphase, n = 7 roots and 19 cells for telophase, n = 8 roots and 28 cells for cytokinesis, and n = 10 roots and 41 cells for postmitotic cells; *tdm3-1*, n = 6 roots and 15 cells for interphase, n = 6 roots and 20 cells for prometaphase–anaphase, n = 6 roots and 20 cells for telophase, n = 6 roots and 25 cells for cytokinesis, and n = 11 roots and 52 cells for postmitotic cells **i,** Representative images of root meristems expressing *pCOILIN::COILIN:RFP* followed by cordycepin treatment and untreated control. **j,** Box plot with jittered points showing quantification of CBs volume in WT and *tdm3-1* after cordycepin treatment and untreated control. On average, CB volume was reduced by 42.7% in WT but by only 18.3% in *tdm3-1*. WT, n = 106 CBs before and n = 91 CBs after cordycepin treatment; tdm3-1, n = 196 CBs before and n = 294 CBs after cordycepin treatment. Quantification was performed using CBs pooled from 3–5 independent roots per condition. **k,** Localization of COILIN:RFP and U2AF35B:RFP CB markers in transiently transfected mesophyll protoplasts. Protoplasts were imaged 16 hrs after transfection. **l,** Representative images and quantification of COILIN:RFP FRAP signal CBs of WT, *tdm3-1* and *35S::TDM3:YFP* plants. Chart shows recovery curves with SDs. WT, n = 8 CBs; *tdm3*, n = 9 CBs; and *35S::TDM3:YFP*, n = 5 CBs. Scale bars: 5 μm (**a,e,k**), 2 μm (**b,c,f**), 1 μm (**l**) and 20 μm (**i**). Scale bar as indicated in (**g**).

In animals, CBs disassemble during mitosis and reassemble in newly formed nuclei at the end of cell division ^36,37^. The transient association of TDM3 with CBs during cytokinesis, together with its interaction with COILIN, suggested that TDM3 may function in CB reassembly at the end of mitosis. To test this hypothesis, we examined CB dynamics in dividing root cells using a line co-expressing COILIN:RFP and the histone marker HTA:YFP (Fig. 2f; Video 3). In wild-type interphase cells, we observed an average of five CBs per nucleus (Fig. 2h). Upon mitotic entry and nuclear envelope breakdown, CBs did not dissolve but instead fused into one to four speckles that persisted in the cytoplasm throughout metaphase and anaphase. CBs reappeared in the newly formed nuclei during telophase, and their number increased to up to ten per nucleus during cytokinesis. In post-mitotic cells of the root transition zone, where cells no longer divide, the number of CBs decreased again to approximately five per nucleus (Fig. 2f-h). Thus, unlike in animals, Arabidopsis CBs do not fully disassemble during mitosis, but instead undergo dynamic reorganization.

Loss of TDM3 reduced the number of CBs in mitotic cells, with the strongest effect observed during cytokinesis, when TDM3 associates with CBs (Fig. f-h). Notably, the absence of TDM3 also decreased CB number in post-mitotic cells of the root transition zone, where TDM3 is not expressed (Fig. 2g,h). In addition, CBs in these post-mitotic cells were larger and less sensitive to dissolution by cordycepin (Fig. 2i,j), a transcriptional inhibitor that disrupts the supply of RNA components required to scaffold CB assembly ^38^. We also observed altered CB behavior in protoplasts derived from terminally differentiated mesophyll cells, which do not express TDM3 (Fig. 2k). CB-associated proteins transiently expressed in transfected protoplasts readily incorporated into CBs in the wild type, but failed to do so in *tdm3* mutants (Fig. 2k). These observations suggest that CBs in *tdm3* mutants are more stable and less dynamic. To further assess CB dynamics in post-mitotic cells, we analyzed COILIN mobility using fluorescence recovery after photobleaching (FRAP). Consistent with observations in animals ^39^, COILIN exhibited low mobility, and its signal showed little or no recovery within the examined time interval in either wild-type or *tdm3* cells (Fig. 2l). However, COILIN mobility increased in post-mitotic cells from plants ectopically overexpressing *TDM3* from a constitutive promoter (Fig. 2l; Extended Data Fig. 7) suggesting that TDM3 enhances the dynamic behavior of CBs.

Together, these results indicate that although TDM3 associates with CBs only transiently at the end of mitosis, it plays an important role in proper CB assembly and has lasting effects on CB architecture and dynamics in post-mitotic cells.

### TDM3 promotes transition from mitotic divisions to differentiation

The reduced stature of *tdm3* mutants suggested impaired growth (Fig. 1b), which was further supported by slower root growth (Fig. 3a and Extended Data Fig. 8). Although *coilin* mutants showed no obvious developmental defects, their root growth was also reduced relative to wild type, albeit less severely than in *tdm3* (Fig. 3a and Extended Data Fig 8). Combined *tdm3 coilin* mutants exhibited severe stunting and infertility (Extended Data Fig. 9), indicating partial functional redundancy of these CB-associated proteins in plant development.

**Fig. 3:**
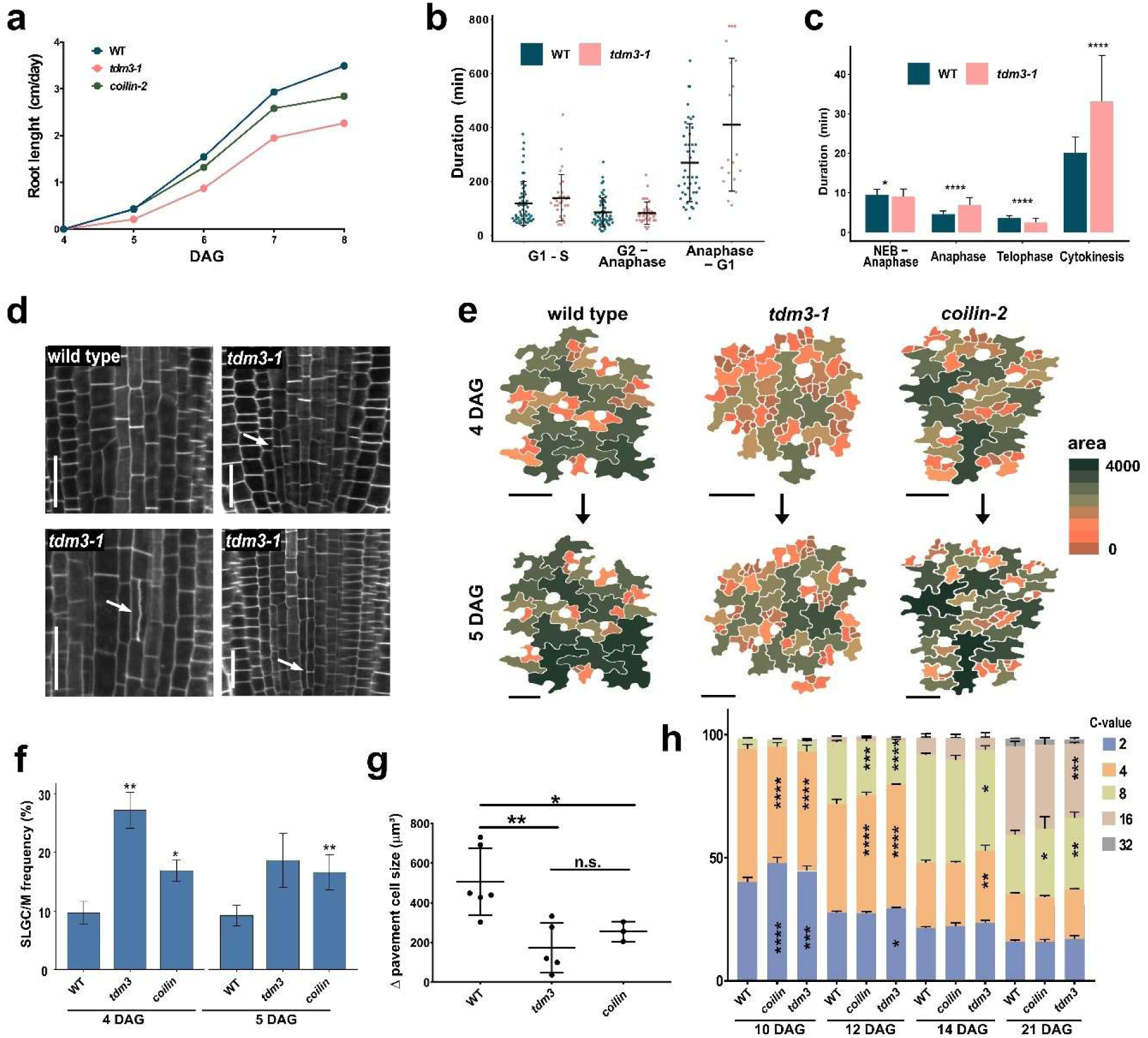
Role of TDM3 in cell division and differentiation. a, Quantification of root growth progression in seedlings from 4 days after germination (DAG) until 8 DAG. **b,** Box plot with jittered points showing quantification of cell-cycle phase duration assessed using the PlaCCI reporter in root meristems. Cohen’s d/Hedges’ g effect sizes indicate the magnitude of differences between genotypes: Anaphase–G1, 0.48/0.47 (medium); G1–S, 0.24/0.24 (small); G2–anaphase, −0.06/−0.06 (negligible). **c,** Quantification of mitotic stage duration using HTA:RFP and TUB4:RFP markers in WT and *tdm3-1*. *(*P < 0.05, ****P < 0.0001*; P-values were determined by Two-sided Wilcoxon rank-sum tests followed by Holm correction for multiple comparisons; for WT n = 89 for NEB–anaphase, 114 cells for anaphase, 36 telophase and 42 for cytokinesis; for *tdm3-1*, n = 84 cells for NEB–anaphase, 114 for anaphase, 32 telophase and 38 cytokinesis). **d,** SR2200 stained root meristems from 5 days old seedlings. Ectopic cell divisions are indicated by arrows. Scale bar = 20 µm. **e,** Spatial maps depicting the epidermal cells in cotyledons in the same area at 4 and 5 DAG. Heat map indicates area of pavement cells and stomatal precursor cells in µm. Scale bar = 50 µm. **f,** Bar plot showing frequency of SLGC and meristemoid cells in cotyledon epidermis *(*P < 0.05, **P < 0.01*; Mann-Whitney U test) **g,** Average changes in pavement cell size between 4 and 5 DAG *(*P < 0.05, **P < 0.01*; Welch ANOVA test). **h,** Nuclear DNA ploidy distribution in true leaves 1 and 2 at different developmental stages *(*P < 0.05, **P < 0.01, ***P < 0.001, ****P < 0.0001*, ANOVA with Šidák correction test; n = 3).

Considering the cell cycle-dependent regulation of TDM3, we analyzed the impact of TDM3 inactivation on cell cycle progression. By tracking the fluorescence cell cycle marker PlaCCI ^40^ in growing roots, we observed that *tdm3* mutants exhibit an extension of the cell cycle phase spanning from cyclin B degradation during anaphase to the onset of CDT1 synthesis in G1 (Video 4, Fig. 3b). Root meristems of *tdm3* mutants also displayed a higher mitotic index (4.09 ± 0.48 in wild type vs. 5.41 ± 1.14 in *tdm3*). A more detailed analysis focused on mitotic progression using chromatin and microtubule markers revealed a substantial prolongation of cytokinesis, the stage that coincides with TDM3 localization in CBs (Videos 5-8, Fig. 2c, Extended Data Fig. 10). This observation indicates that TDM3-mediated co-assembly with CBs facilitates completion of mitosis.

In *tdm3* mutants, we also noticed ectopic cell divisions within root cell, where a single cell appeared to undergo an additional division, giving rise to two smaller daughter cells (Fig. 3d). This phenotype suggests impaired termination of mitotic activity and a delayed transition to differentiation. However, further analysis in roots was complicated by the variability of stereotypic root cell division patterns. To investigate this phenomenon in a more tractable system, we examined the development of leaf epidermal cells, which enables the tracking of precursor cell divisions and their differentiation into stomatal and pavement cells across the surface of young leaves (Fig. 3e)^41^. These precursor cells also express TDM3 in CBs (Extended Data Fig. 3d).

The epidermis of developing cotyledons in *coilin* and *tdm3* mutants contained a higher proportion of undifferentiated precursor cells than wild-type seedlings at a comparable growth stage (Fig. 3e,f). Tracking epidermal cells in developing cotyledons between 4 and 5 days after germination (DAG) further revealed higher rates of cell proliferation and lower rates of cell enlargement in *tdm3* and *coilin* mutants relative to the wild type (Figs. 3e,g and Extended Data Figs. 11–13). These observations suggest prolonged amplification cycles and delayed differentiation of meristemoid and stomatal lineage ground cells (SLGC) into stomata and large pavement cells. A similar trend was observed in true leaves, where reconstruction of cell divisions in developing stomatal complexes tended to show an increase in spacing divisions of stomatal lineage ground cells in *tdm3* plants (Extended Data Figs. 14 - 16). In Arabidopsis, the transition from amplifying cell divisions to pavement cell differentiation is accompanied by endoreplication and cell enlargement ^42,43^. Consistent with a delayed transition to differentiation, developing true leaves of *tdm3* and *coilin* mutants showed a slower accumulation of higher-ploidy cells (Fig. 3h). Together, these data suggest that TDM3 and COILIN promote the transition from proliferative mitotic divisions to terminal cell differentiation.

### TDM3 impacts pre-mRNA splicing

COILIN broadly influences gene expression and splicing across multiple model organisms, including Arabidopsis ^25,44–47^. Given the association of TDM3 with COILIN and CBs, we profiled transcriptomes of 4-days-old seedlings in *coilin* and *tdm3*. While *colin* mutants displayed 2,799 differentially expressed genes (DEGs), only 147 DEGs were identified in *tdm3* (Fig.4a, Supplementary Table 1). Notably, a substantial portion *of tdm3* DEGs overlapped with those in *coilin*. These shared genes were concordantly regulated between the mutants (Fig. 4b and Extended Data Fig. 17). To identify broader expression patterns, we performed unsupervised clustering of all 2,855 DEGs from both mutants (Extended Data Fig. 18). Cluster 3, which contains DEGs showing strongest upregulation in *tdm3* and a weaker upregulation in *coilin*, was enriched for mitotic and cell cycle functions. This pattern is consistent with an increased proportion of mitotic cells, which is more pronounced in *tdm3* compared to *coilin*.

**Fig. 4:**
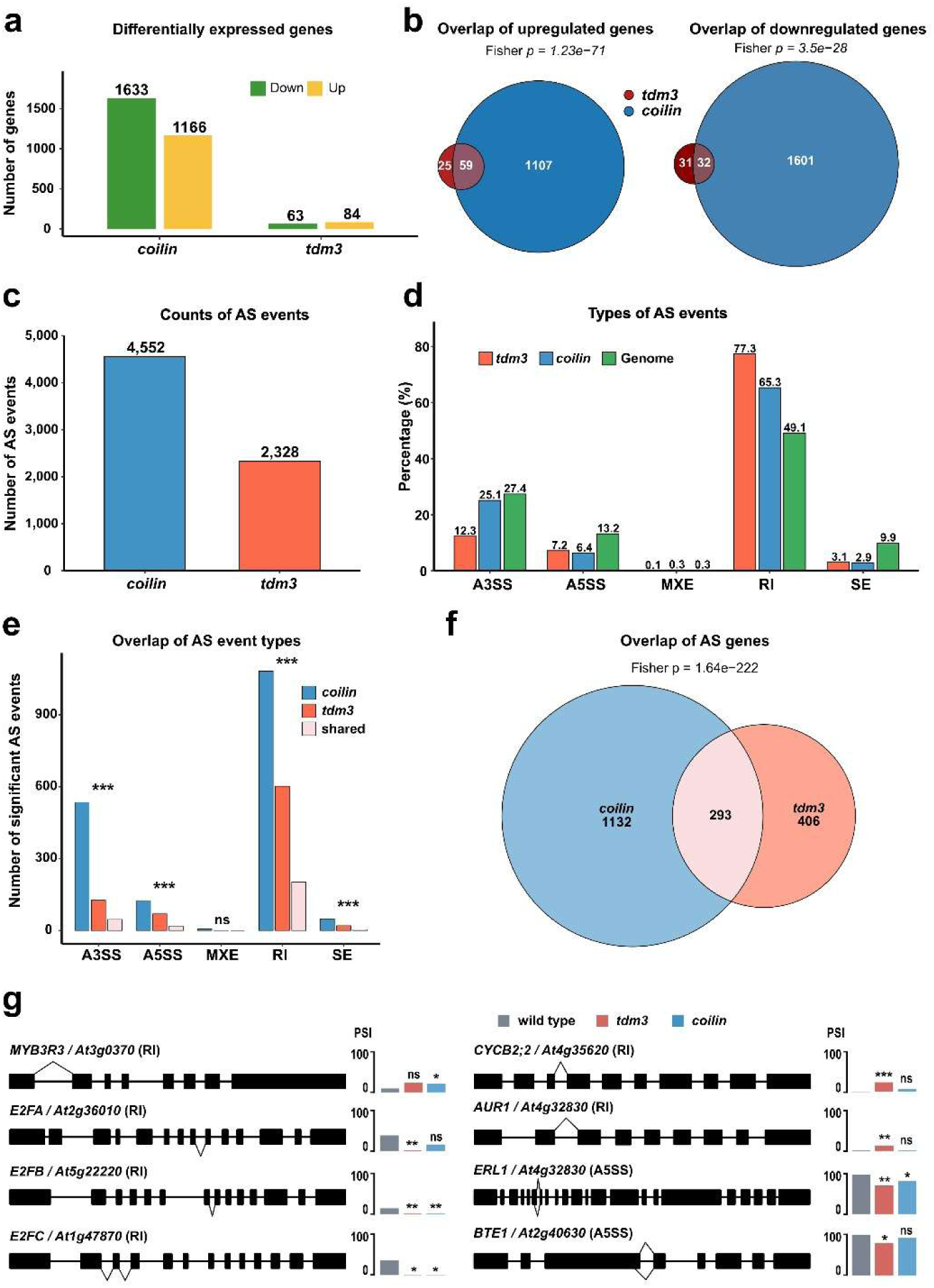
Gene expression and splicing analyses. a, Bar chart showing DEGs in *tdm3* and *coilin* relative to WT (adjusted *P < 0.05*). **b,** Area-proportional Venn diagrams showing overlap between upregulated (left) and downregulated (right) DEGs in *tdm3-1* and *coilin-2*. Significance was assessed by Fisher’s exact test. **c,** Total numbers of significant alternative splicing (AS) events detected in *tdm3* and *coilin* (FDR < 0.05). **d,** Distribution of AS event classes relative to the AtRTD3 reference transcriptome: Alternative 3′ Splice Site (A3SS), Alternative 5′ Splice Site (A5SS), Mutually Exclusive Exons (MXE), Retaiend Introns (RI) and Skipped Exons (SE). Retained intron events are enriched in both *coilin* and *tdm3*. **e,** Bar chart showing numbers of significant AS events across different classes in *coilin* and *tdm3*, and of the shared events. Asterisks indicate significance of the overlap determined by Fisher’s exact test *(***P < 0.001*; ns, not significant). **f,** Area-proportional Euler diagram showing overlap between genes affected by significant AS events (ΔPSI > 0.1, FDR < 0.05) in *tdm3-1* and *coilin-2*. The shared set of 293 genes indicates extensive convergence of splicing defects between the two mutants (*P = 1.64 × 10^-222^*, Fisher’s exact test**). g,** Examples of AS genes involved in cell cycle regulation and differentiation. Splicing events enriched in WT and *tdm3* or *coilin* mutants are indicated above and below each gene model, respectively. PSI (percent spliced in) values for the indicated events are shown in the charts to the left of the gene models.

Whereas TDM3 had only a modest effect on gene expression, its inactivation strongly impacted pre-mRNA splicing. A total of 4,552 significant alternative splicing (AS) events were detected in *coilin* and 2,328 in *tdm3* (Fig. 4c and Extended Data Fig. 19, Supplementary Table 2,3). In both mutants, intron retention (IR) was the predominant event type, followed by alternative 3′ splice sites (A3SS). Although *coilin* displayed a higher total number of AS events compared to *tdm3*, the distribution of these events slightly differed between the mutants (Fig. 4d). While *coilin* exhibited a higher absolute number of IR events (2,973 vs. 1,800 in t*dm3*), IR accounted for a greater proportion of total AS events in *tdm3* (77% vs. 65% in coilin). This predominant effect on intron retention events indicates that TDM3 primarily influences splicing efficiency rather than splice-site selection ^17,48^.

To define high-confidence AS events, we applied stringent filtering criteria requiring at least 10 reads per junction and a minimum ΔPSI of 10%, yielding 1,799 events in *coilin* and 810 in *tdm3* (Extended Data Fig. 20). A significant proportion of these events was shared between the mutants (Fig. 4e), suggesting that TDM3 and COILIN target a common subset of pre-mRNA splice sites. These significant AS events were associated with 1,832 genes, 293 of which were shared between *coilin* and *tdm3* plants (Fig. 4f, Supplementary Table 4). Strikingly, genes showing concordant splicing changes in *tdm3* and *coilin* included several components of the DREAM complex (*E2FA, E2FB, E2FC, MYB3R3*, and *BTE1*), which plays a fundamental role in cell fate transition through acting as a master regulator of cell cycle quiescence and terminal differentiation^49–53^. We also identified *ERL1*, a receptor-like kinase involved in epidermal cell differentiation, as well as the mitotic factors *CYCB2;2* and *AUR1* (Fig. 4g) ^54,55^. The shared splicing events affecting key regulators of cell proliferation and differentiation imply a molecular link between CB-associated RNA processing and developmental cell fate transitions.

## Discussion

In contrast to animal cells, where CBs disassemble upon nuclear envelope breakdown ^36,37^, plant CBs persist as large cytoplasmic condensates throughout mitosis and undergo extensive remodeling following their incorporation into newly formed daughter nuclei (Fig. 2f-h). Our results indicate that TDM3 facilitates this remodeling process. Although TDM3 localizes to CBs only transiently during cytokinesis, this interaction has lasting effects on CB properties in post-mitotic cells. CBs continuously exchange components with the surrounding nucleoplasm ^24,39^, and this dynamic behavior is reduced in *tdm3* mutants but enhanced in plants ectopically overexpressing TDM3 (Fig. 2i–l). These findings suggest that TDM3 acts as mitosis-associated CB-assembly factor that promotes the formation of dynamic condensates by preventing interactions that would otherwise give rise to more rigid structures. Similarly, in human cells, CB assembly depends on COILIN oligomerization through its N-terminal domain and on interactions with Nopp140, which prevent the formation of aberrant COILIN fibrils ^56^. Together, these findings suggest that both plant and animal cells employ dedicated mechanisms to ensure the proper assembly and material properties of CBs.

CBs promote efficient pre-mRNA splicing by regulating the supply, quality control, and modification of the core splicing machinery ^21–23^. Our findings suggest that CB function contributes to the transition from cell proliferation to differentiation. This role may partly reflect the increased demand for RNA processing during cellular reprogramming. However, transcriptome analyses also revealed splicing defects in several key regulators of cell cycle progression and cell fate determination (Fig. 4g). Although COILIN exerts a broader effect on splicing, these regulatory transcripts were more strongly affected in *tdm3* than in *coilin* mutants. Consistent with this observation, *tdm3* mutants display more severe developmental phenotypes than *coilin* mutants, and combined mutations further exacerbate these defects (Figs. 1a and 3 and Extended Data Fig. 9). These observations suggest that TDM3 is not merely a structural regulator of CB assembly but may additionally contribute to the selective processing of transcripts that govern cell cycle regulation and differentiation.

Our previous work revealed that the closely related TDM1 protein also regulates a major developmental transition by promoting the termination of meiosis and the onset of microspore differentiation ^7^. Although both proteins act through RNP condensates, they appear to do so via distinct mechanisms: TDM1 regulates translation through cytoplasmic P-bodies, whereas TDM3 promotes Cajal body assembly and affects alternative splicing. Together, these findings suggest that the TDM protein family has evolved as a versatile regulator of developmental transitions by modulating the behavior of distinct RNP condensates. More broadly, our work supports a model in which dynamic reorganization of RNP condensates represents a general mechanism for coupling cell-cycle transitions to the establishment of new cellular identities.

## Material and methods

### Plant material and growth conditions

*Arabidopsis thaliana* seeds were surface-sterilised and grown in soil under controlled-chamber conditions at 21 °C, 51–60% relative humidity, and a 16/8 or 12/12-hour light/dark photoperiod. The Columbia-0 (Col-0) ecotype was used as the wild-type (WT) reference. The *tdm3-1* (GK-17306), and *coilin-2* (SALK_148589) ^46^ lines were obtained from the Nottingham Arabidopsis Stock Centre (NASC). The *ccs52a1-1* (SALK_083656) has been previously described ^33,57^. Genotyping of all lines was performed using allele-specific primers (Supplementary Table 5).

### Phenotyping assays in roots and flow cytometry

For phenotypic analysis, *Arabidopsis thaliana* seeds were surface-sterilised and sown on half-strength Murashige and Skoog (½ MS) medium containing of 2.45 g/L MS salts with vitamins and MES buffer (Duchefa), solidified with 7 g/L agarose. Media were prepared either with or without 10 g/L sucrose (designated as MS⁺ and MS⁻, respectively). Plates were positioned vertically and incubated in a growth chamber at 21 °C under a 16/8-hour light/dark photoperiod with LED illumination. Root growth assays were quantified as cumulative primary root growth relative to 4 days after germination (DAG, 5–4, 6–4, 7– 4 and 8–4 DAG intervals). Digital images of the seedlings were acquired daily using a flatbed scanner or digital imaging system. Root growth was quantified in Fiji/ImageJ ^58^ by manually tracing the primary root length. For the flow cytometry analysis, WT, *tdm3-1* and *coilin-2* mutants were grown in soil (Jiffy Pellets 7C) under long-day conditions (16 h light/8 h dark, Lumilux Cool White lm, 50 to 70 μmol m−2 s−1) at 21 °C. After 10, 12, 14 and 21 days of growth, the first pair of true leaves (Day 10) or the second true leaf only (Day 12, 14 and 21) was harvested and flash-frozen in liquid nitrogen. Frozen leaf material was chopped on a petri dish with a razor blade in 200 μl Nuclei Extraction Buffer (Cystain UV Precise P, Sysmex) and 800 μl Staining Buffer (Cystain UV Precise P, Sysmex) was added to the extracts. The homogenate was filtered on a 30 μm Cell Trics filter. Measurements of the DNA content were performed with QA Sheath Fluid (Quantum Analysis, Germany) on a Quantum P (Model QA FCM QP 2, Quantum Analysis GmbH, Germany) flow cytometer excited by illumination at 395 nm. The ploidy levels (2C, 4C, 8C, 16C and 32C fractions) were determined from DAPI fluorescence measurements using Floreada (https://floreada.io). The relative cell population (2C, 4C, 8C, 16C and 32C) sizes, calculated as the percentage of all cells, averaged across three biological replicates.

### Transient expression in protoplasts

The TDM3:YFP construct was previously generated by cloning the TDM3 coding sequence into the Gateway destination vector pGWB441^7,59^. The following subnuclear RFP-fused marker constructs were generated: COILIN:RFP for Cajal bodies ^34^, HYL1:RFP for Dicing bodies ^60^, U2AF35B:RFP for splicing speckles (nuclear splicing) ^61^ and FIB1:RFP for the nucleolus ^62^. The corresponding cDNAs (COILIN, HYL1, U2AF35B, and FIB1) were amplified using gene-specific primers (Supplementary Table 5). Entry clones were generated in pENTR™/D-TOPO and subsequently recombined into pGWB460 to generate RFP-tagged expression vectors. For BiFC (Bimolecular Fluorescence Complementation) assays, entry clones were recombined with BiFC destination vectors pGWcY and pnYGW ^59^. The following combinations were used: pGW-TDM3-cY and pnY-COILIN-GW, to assess protein–protein interactions involving TDM3 and COILIN.

The Arabidopsis mesophyll protoplast transfection protocol was performed as previously described ^7^. After transformation, protoplasts were incubated in W5 solution for 16 h at room temperature and imaged using a Zeiss LSM780 microscope, and image processing was carried out with ZEN software.

### Plasmid construction and generation of transgenic Arabidopsis lines

All primers used in this study are listed in Supplementary Table 5. To generate the *pTDM3::TDM3:YFP* reporter lines, a genomic fragment encompassing the TDM3 promoter and open reading frame (ORF) was amplified using the primers TDM3fw_TOPO and TDM3rev_TOPO. The resulting PCR product was cloned into the pENTR™/D-TOPO vector (Invitrogen), followed by LR recombination into the destination vector pGWB640 (YFP C-terminal fusion)^59^. The construct was introduced into homozygous *tdm3-1* mutant plants using Agrobacterium-mediated floral dip transformation ^63^. For the *pCOILIN::COILIN:RFP* reporter lines, a genomic fragment including the COILIN promoter and ORF was amplified using COILINfw_prom_TOPO and COILINrev_prom_TOPO, cloned into pENTR™/D-TOPO, and recombined into pGWB659 (RFP C-terminal fusion). The resulting construct was transformed into homozygous *coilin-2* and *tdm3-1* plants via floral dip. To generate the *pHTA10::HTA10:eYFP* line (HTA:YFP), the promoter and genomic coding sequence of HTA10 were amplified using HTAYFP_fw and HTAYFP_rev, and the product was cloned into pGWB640 for C-terminal eYFP fusion. The construct was transformed into WT plants via floral dip. For constitutive overexpression, the *35S::TDM3-YFP* construct was generated by amplifying the TDM3 cDNA using At3G51280_FW and AT3G51280_nostop_R. The amplicon was cloned into pENTR™/D-TOPO and recombined into pGWB641(YFP-tag under the 35S promoter). The construct was introduced into *tdm3-1* homozygous plants via floral dip. For all reporter constructs, non-segregating homozygous T3 or higher-generation lines were selected and used for phenotypic and microscopic analyses. Additionally, the following published reporter lines were used for imaging root tissues and cellular structures: *HTA10:RFP* (chromatin visualization, HTA:RFP) ^64^, *TagRFP:TUB4* (microtubule and mitotic spindle marker) ^65^, *PlaCCI* (cell cycle phase indicator) ^40^, and *PCNA:TagRFP* (replication complex)^64^. Transgenic reporter lines were crossed with desired mutants or other reporter lines. F1 progenies were screened by fluorescence microscopy, and homozygous, non-segregating F3 lines were selected for downstream imaging and analyses.

### Analysis of leaf epidermal cells

For the pavement cell division analysis, plants were stratified in darkness for three days and subsequently grown at 21°C. The abaxial epidermis of the cotyledons was imprinted using polyvinyl impression material (3M Light Body Fast Set Std, ref. 7301), and the resulting replicas were covered with nail polish. The nail polish was examined by scanning electron microscopy (TM-1000, Hitachi). On 4 to 5 DAG, 3–5 cotyledons were monitored. Quantification focused on pavement cells and stomatal precursor cells (meristemoids and stomatal lineage ground cells), including their division activity, area changes, and developmental progression. Approximately 50 cells per region were scored, and cell-tracking analysis was performed automatically using R by retrieving progeny cells based on cell surface overlap between consecutive time points.

For the epidermal cell analysis in true leaves, WT and *tdm3-1* plants were grown in soil (Jiffy Pellets 7C) under long-day conditions (16 h light/8 h dark, Lumilux Cool White lm, 50–70 μmol m⁻² s⁻¹) at 21 °C. After 10 and 21 days of growth, the abaxial surface of the second true leaf (Leaf #2) was analysed from the imprints. From the imprint images, cellular complexes of developing stomata were manually tracked. This included identification of meristemoids and stomatal lineage ground cells. Per genotype, approximately twenty stomatal complexes were analysed in each of three independent leaves (total n = 60 per genotype). For cell tracking, daughter cells at Day 21 were assigned based on surface overlap with mother cells at Day 10.

### Cytology and confocal microscopy

Plant tissues expressing protein markers were imaged using a Zeiss LSM780 (C-Apochromat 40×/1.2 W), with optional Propidium-iodide or DAPI staining. Analysis of ovules was performed as previously described ^66^. Samples were mounted in VECTASHIELD® and imaged using a LSM700 (Plan-Apochromat 63x/1.4, Oil) and LSM880 with Airyscan (Plan-Apochromat 63x/1.4, Oil).

For quantification of Cajal bodies during the cell cycle, roots expressing *pCOILIN::COILIN:RFP* were fixed in formaldehyde, washed in PBS, stained with DAPI for 1 h and mounted in VECTASHIELD under a coverslip. Samples were imaged using a Zeiss LSM780 confocal microscope equipped with a C-Apochromat 40×/1.2 W objective. Cell-cycle stages were assigned based on chromatin morphology and Cajal body numbers were counted manually.

For chemical treatments, 4–5 DAG seedlings were incubated in MS medium containing 50 µM MG132 or 0.6 mM cordycepin for 1 hour. Imaging before and after treatment was performed using a Zeiss LSM780 (C-Apochromat 40×/1.2 W). Cajal bodies in cordycepin-treated plant tissues were segmented using the surface creation algorithm in Imaris v9.2.1 (Bitplane AG). Segmentation parameters were manually adjusted to optimize detection, and the resulting surface volumes were exported for quantitative analysis. FRAP experiments were performed using an LSM780 confocal microscope (Zeiss). Regions of interest were photobleached using a 514 nm laser at 100% intensity for YFP and a 661 nm laser at 100% intensity for TagRFP. Fluorescence recovery was monitored by time-lapse imaging for 180 s.

### Live cell imaging of root cells

For analysis of mitosis, seedlings were grown vertically on ½ MS plates for 4 days, then transferred to round microscopy chambers with agar overlay. Time-lapse imaging was performed on a Zeiss LSM780 (Plan-Apochromat AIR 20×/0.8) for up to 1 hour, acquiring Z-stacks every 1 minute. Timing of mitotic stages was quantified manually in ZEN Black and Fiji/ImageJ ^58^. HTA10:RFP and TUB4:TagRFP markers were used to define duration of individual mitotic stages.

Long-term live imaging of growing roots was performed on a plant-optimised vertical stage Zeiss LSM880 with Airyscan detector (Carl Zeiss AG). Root tracking was based on the Matlab object tracking macro developed based on Wangenheim et al. ^67^. 4-5 DAG seedling was placed in the coverglass chamber, and its root tip was imaged using Plan-Apochromat 20×/0.8 objective; the Z-stacks encompassing at least half of the root diameter were acquired at 5-to-15-minute intervals for 12–24 hours with an Airyscan detector (CFP ex: 458 nm; YFP ex: 514 nm, both em: BP 465-505 + LP 525 Airyscan; mCherry ex: 561nm, em: BP 495-550 + LP 570 Airyscan). Airyscan processing was performed using ZEN black 2.6 software. Time stitching of z-section and, z- and time alignment were performed in the ZEN Blue 3.4 software (Carl Zeiss AG) and analysed in Arivis4D (Zeiss, Arivis AG). Datasets were first aligned to individual nuclei, then segmented for each fluorescent channel and automated bleaching correction was applied. After initial processing, nuclei were manually curated for accurate tracking, timing, signal evaluation, and correction of segmentation errors. For quantification, temporal profiles of fluorescent signal were used to assign cell cycle stages based on the expression PlaCCI markers ^40^.

For *pCOILIN::COILIN:RFP* × *pHTA10::HTA10:YFP* crosses, enabling simultaneous visualization of Cajal bodies and chromatin, live-cell time-lapse imaging was performed on a Zeiss LSM780 confocal microscope equipped with a Plan-Apochromat AIR 20×/0.8 objective. Vertically grown seedlings (4–5 DAG) were imaged every 1.5–2 minutes over the course of 1 hour. Datasets were processed using ZEN software and Imaris. For the analysis of *pCOILIN::COILIN:RFP* with *pTDM3::TDM3:YFP* lines, used to study the spatiotemporal relationships between Cajal bodies and cell cycle progression in root meristem cells, 4–5 DAG seedlings were imaged in a horizontal configuration using the Zeiss LSM880 with Airyscan module. Roots were tracked live using a Plan-Apochromat 20×/0.8 DIC M27 objective, acquiring Z-stacks encompassing at least half of the root diameter at 5–15-minute intervals for 12–24 hours. Datasets were processed using ZEN software and further analysed in Arivis 4D 4.1.1.

### Yeast two-hybrid assay

Yeast two-hybrid (Y2H) experiments were conducted using the Matchmaker™ GAL4-based two-hybrid system (Clontech). For pairwise interaction assays, bait and prey vectors were transformed into Y187 and AH109 yeast strains, respectively. Protein-protein interactions were assessed using the mating method. Yeast cells carrying both plasmids were initially selected on SD-Leu-Trp medium, and positive interactions were identified on SD-Leu-Trp-His selection media with an appropriate concentration of 3AT (3-amino-1,2,4-triazole). Empty vectors were included as negative controls to rule out auto-activation. For Y2H constructs, TDM3, TDM1 and TDM2 cDNAs were amplified using the primers At3G51280_fw_Nde, At3G51280_rev_Sal, TDM1_fw_Nde, TDM1_Rev_SaI, AT5G44330_FW_Nde and AT5G44330_Rev_Sal. The resulting fragments were digested with NdeI and SalI restriction enzymes and cloned into the pGBKT7 (bait) vector. For COILIN, CCS52A1, APC7, and CDC27(HOBBIT), cDNA was amplified using the primers: COILINfw_TOPO, COILINfull_rev, AT2G20000_fw, AT2G20000_rev, APC7.TOPO.F, APC7.TOPO.STOP.R, CCS52A.1.TOPO.F and CCS52A.1.TOPO.STOP.R. The amplified products were first cloned into the pENTR™/D-TOPO vector, and destination vectors containing the respective cDNAs were subsequently generated through LR recombination, resulting in constructs in the pGADT7-DEST (prey) plasmid.

### Phylogenetic analysis

Protein sequences (Supplementary Table 6) were retrieved using NCBI BLAST ^68^ and PANTHER ^69^ databases, and multiple sequence alignment was performed using MUSCLE ^70^. Phylogenetic analysis was performed using the Maximum Likelihood method with the Jones–Taylor–Thornton (JTT) amino acid substitution model ^71^. Branch support was assessed using the bootstrap method with 1,000 replicates. A uniform rate among sites was assumed. Positions with less than 95% site coverage were excluded using partial deletion. The initial tree was generated automatically, and tree topology was optimized using the Nearest-Neighbor-Interchange (NNI) heuristic method. The analysis was done in MEGA11 ^72^. A full list of species and accession numbers used is provided in Supplementary Table 5.

### Statistical analysis

All statistical analyses were performed using GraphPad Prism v.7 (GraphPad Software) and RStudio v.1.3.1073 (Posit PBC). Statistical tests, sample sizes, and exact n values are indicated in the corresponding figure legends. Two-sided statistical tests were used throughout the study, and multiple-comparison corrections were applied as indicated. Root growth trends were additionally quantified using area under the curve (AUC) analysis integrating cumulative root growth across the measured developmental time course. Flow-cytometry datasets were analyzed using ANOVA followed by Šídák multiple-comparison correction ^73^. Effect size analysis (Cohen’s d and Hedges’ g) ^74^ was additionally performed for PlaCCI-derived cell-cycle phase duration datasets to assess biological trends independent of statistical significance. Cell-cycle progression datasets derived from HTA10/TUB4 markers and Cajal body quantifications at individual mitotic stages were analysed using two-sided Wilcoxon rank-sum tests ^75^ followed by Holm correction ^76^ for multiple comparisons.

Stomatal lineage analysis was performed using epidermal cotyledon imprints obtained at 4 and 5 DAG. ΔPC area was calculated as the cell area difference between Day 5 and Day 4 (ΔPC = PCDay5 − PCDay4). Because several datasets showed unequal variances and limited sample sizes, Welch-corrected statistical approaches were used where appropriate, including Welch’s ANOVA followed by pairwise Welch comparisons ^77^. Stomatal lineage analyses were performed using cotyledon epidermal imprints from the 4–5 DAG time-course experiment together with additional independent cotyledon imprints collected at the same developmental stages. These additional samples were not included in temporal analyses but were used for quantitative phenotyping. Non-stomatal epidermal cells were operationally classified into size-defined categories based on cell area. Cells up to 90 μm² were classified as M/SLGC-like cells ^41^, whereas cells larger than 90 μm² were classified as pavement cells (PCs). Frequencies of individual cell classes were calculated per biological replicate as proportions of the total quantified epidermal population. Individual stomatal lineage ground cells (SLGCs) were manually identified and scored in each biological replicate, and the proportion of dividing SLGCs was calculated for each sample. Differences in SLGC division frequency between WT and each mutant genotype were assessed using the Wilcoxon rank-sum test ^75^. Statistical significance is indicated in graphs as follows: *P < 0.05, **P < 0.01, ***P < 0.001, and **** P < 0.0001.

### Analysis of protein structures

Protein structures of TDMs family were retrieved from the Protein Data Bank (PDB) and analysed using UCSF ChimeraX ^78^ for visualization and structural comparison (AF-Q9SUC3-F1-v4 for TDM1, AF-Q9FKV5-F1-v4 for TDM2, AF-Q9SD20-F1-model_v4 for TDM3, AF-Q8GXU5-F1-v4 for SDI1, and AF-Q8L730-F1-v4 for SDI2) and were modelled using AlphaFold3 ^79^ and analysed in ChimeraX. D-box (destruction box) motif in the TDM3 sequence VEAEDAYRRA (positions 194–197) was identified using GPS-ARM ^80^. Conservation profiles of TDM proteins were generated from multiple sequence alignments using Shannon entropy-based conservation scoring implemented in R using the Biostrings package ^81^. Sequence conservation was calculated as an inverse Shannon entropy score according to C_i_=H_max_−H_i_, where C_i_ represents conservation at alignment position i, H_i_ corresponds to positional Shannon entropy, and H_max_ represents the maximal entropy observed across the alignment.

### RNA-seq and differential expression analysis

Total RNA was extracted from 1 g of *Arabidopsis thaliana* Col-0, *tdm3-1* and *coilin-2* seedlings using the RNeasy Plant Mini Kit (Qiagen), according to the manufacturer’s instructions. Three independent biological replicates were prepared for each genotype. Genomic DNA was removed by treating 100 µg of total RNA with DNase I and using the TURBO DNA-free Kit (Invitrogen). Strand-specific RNA-seq libraries were prepared from 10 µg of DNase-treated RNA using the TruSeq Stranded Total RNA Kit with Ribo-Zero Plant (Illumina). An additional rRNA depletion step was performed using the FastSelect rRNA Plant Kit (Qiagen). Libraries were sequenced on an Illumina NovaSeq platform to generate 150-bp paired-end reads.

Raw reads were assessed and adapter-trimmed using Trim Galore version 0.6.10 ^82^. Trimmed reads were aligned to the *Arabidopsis thaliana* reference genome TAIR10 using STAR version 2.7.10b ^83^, guided by the AtRTD3 transcript annotation ^84^.

Gene-level counts were generated using featureCounts from Rsubread v2.16.1 ^85^ with the AtRTD3 annotation in paired-end, reverse-stranded mode. Multi-mapping reads and PCR duplicates were excluded. Lowly expressed genes were removed using filterByExpr from edgeR ^86^. Surrogate variables were estimated using svaseq from the sva package ^87^ and included in the DESeq2 model design together with genotype. Differential expression analysis was performed using DESeq2 v1.40.2 with the Wald test, and P values were adjusted using the Benjamini-Hochberg method ^88^. Log2 fold changes were shrunk using apeglm. Genes with an adjusted P value < 0.05 were considered differentially expressed.

For clustering, variance-stabilized counts were generated using DESeq2 and averaged across biological replicates for each genotype. The union of differentially expressed genes was clustered by k-means clustering with k = 8 using ClusterGVis ^89^. Gene Ontology enrichment analysis for each cluster was performed using clusterProfiler ^90^, with all expressed genes as the background set. The top 5 enriched biological process terms were visualized for each cluster.

Differential alternative splicing was analysed using rMATS v4.3.0 ^91^ with the AtRTD3 annotation. Events were classified as skipped exons, alternative 5′ splice sites, alternative 3′ splice sites, mutually exclusive exons or retained introns. Significant alternative splicing events were initially defined by FDR < 0.05. A more stringent high-confidence set was then generated by retaining only events supported by at least 10 junction-spanning reads and showing an absolute change in percent spliced in of at least 10% (|ΔPSI| > 0.1), with FDR < 0.05. The distribution of significant events across splicing classes was compared with the corresponding distribution in the AtRTD3 annotation for statistical significance.

Overlap between significant alternative splicing events in *tdm3-1* and *coilin-2* was assessed by coordinate-based matching within each event class. Statistical significance of overlap was tested using Fisher’s exact test, using all detected events in the corresponding class as the background. Gene-level splicing target sets were generated by assigning significant alternative splicing events to their corresponding genes. Shared and genotype-specific gene sets were visualized using area-proportional Euler diagrams. Gene Ontology enrichment analysis of shared, *tdm3*-specific and *coilin*-specific splicing targets was performed using ShinyGO ^92^, and redundant terms were filtered before visualization.

## Acknowledgements

This work was supported by the Czech Science Foundation (22-31712S). M.D. is a post-doctoral fellow of Flanders Research Foundation (FWO no. 12Q7923N). Microscopy was performed at the Imaging Facility of the IEB AS CR and the core facility of CEITEC Masaryk University CELLIM funded by MEYS CR (LM2023050 Czech-BioImaging). We further acknowledge support of Plant Science Core Facility with plant cultivation.

## Description of videos

**Video 1.** Time-lapse imaging of TDM3:YFP (yellow) and COILIN:RFP (violet) in the Arabidopsis root apical meristem. Images were acquired using a VertiScope microscope. Elapsed time in the HH:MM:SS format and scale bar are indicated.

**Video 2.** Time-lapse imaging of TDM3:YFP (yellow) and COILIN:RFP (violet) throughout the G2 phase to cytokinesis in a single Arabidopsis root apical meristem cell. Images were acquired using a VertiScope microscope. Elapsed time in HH:MM:SS and scale bar are indicated.

**Video 3.** Time-lapse imaging of COILIN:RFP (violet) and HTA:YFP (yellow) in the Arabidopsis root apical meristem. Images were acquired by confocal microscopy. Elapsed time in min and scale bar are indicated.

**Video 4.** 12hrs time-lapse imaging of the PlaCCI cell-cycle reporter in the Arabidopsis root apical meristem, illustrating progression of individual root cells through cell cycle. The reporter consists of CDT1a–CFP (blue), CYCB1;1–YFP (yellow) and HTA10–RFP (red). Images were acquired using a VertiScope microscope. Scale bar as indicated.

**Video 5.** Time-lapse imaging of mitotic progression in root cells in the wild type using HTA10:RFP to visualize chromatin. Images were acquired using a confocal microscope. Elapsed time in min and scale bar are indicated.

**Video 6.** Time-lapse imaging of mitotic progression in root cells in *tdm3-1* using HTA10:RFP. Images were acquired using a confocal microscope. Elapsed time in min and scale bar are indicated.

**Video 7.** Time-lapse imaging of mitotic progression in root meristem in the wild type using TUB4:TagRFP to visualize microtubule dynamics. Images were acquired using a confocal microscope. Elapsed time in min and scale bar are indicated.

**Video 8.** Time-lapse imaging of mitotic progression in root meristem in *tdm3-1* using TUB4:TagRFP to visualize microtubule dynamics. Images were acquired using a confocal microscope. Elapsed time in min and scale bar as indicated.

