## Supplemental Data for "A Cajal body assembly factor regulates cell fate transitions in Arabidopsis"

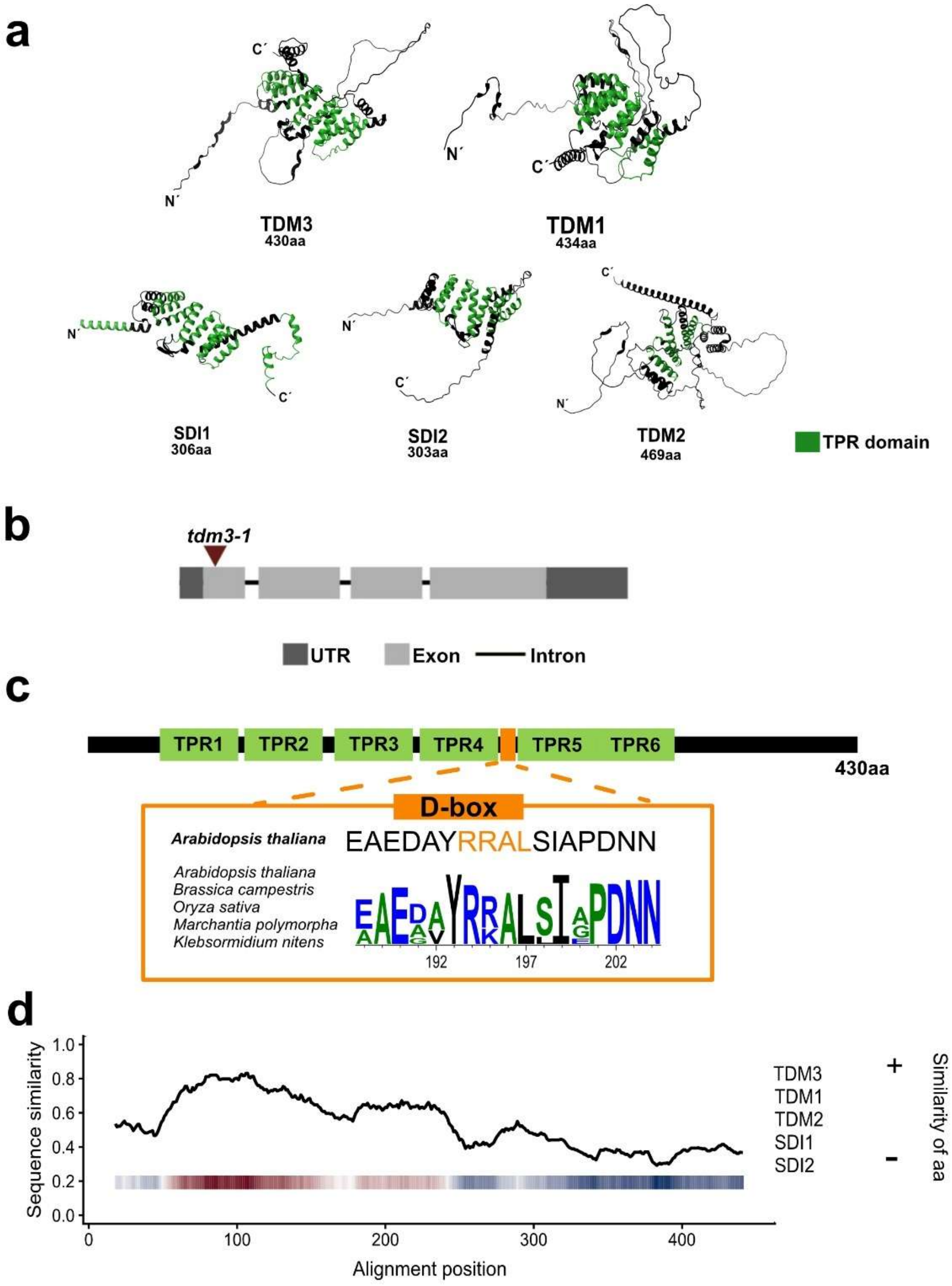


**Extended Data Fig.1 | Genetic and structural characterization of TDM3. a,** AlphaFold-predicted structures of Arabidopsis TDM/SDI proteins illustrating the conserved α-helical organization of TPR domains across the family. TPR domains are highlighted in green. **b,** Schematic representation of the T-DNA insertion site in the *TDM3* gene in the *tdm3-1*; exons, introns, and UTRs are indicated. **c,** Domain organization of TDM3 showing six TPR domains and a conserved D-box motif. Sequence alignment of TDM3 homologs from representative plant species revealed conservation of the D-box region across land plants and *Klebsormidium nitens*. **d,** Sequence conservation analysis of Arabidopsis TDM/SDI proteins based on multiple sequence alignment. Sequence similarity scores across the alignment were calculated using Shannon entropy-derived conservation metrics and visualized together with positional similarity heat maps. The N-terminal region displayed higher sequence conservation, whereas the C-terminal region showed increased sequence divergence among TDM and SDI family members.

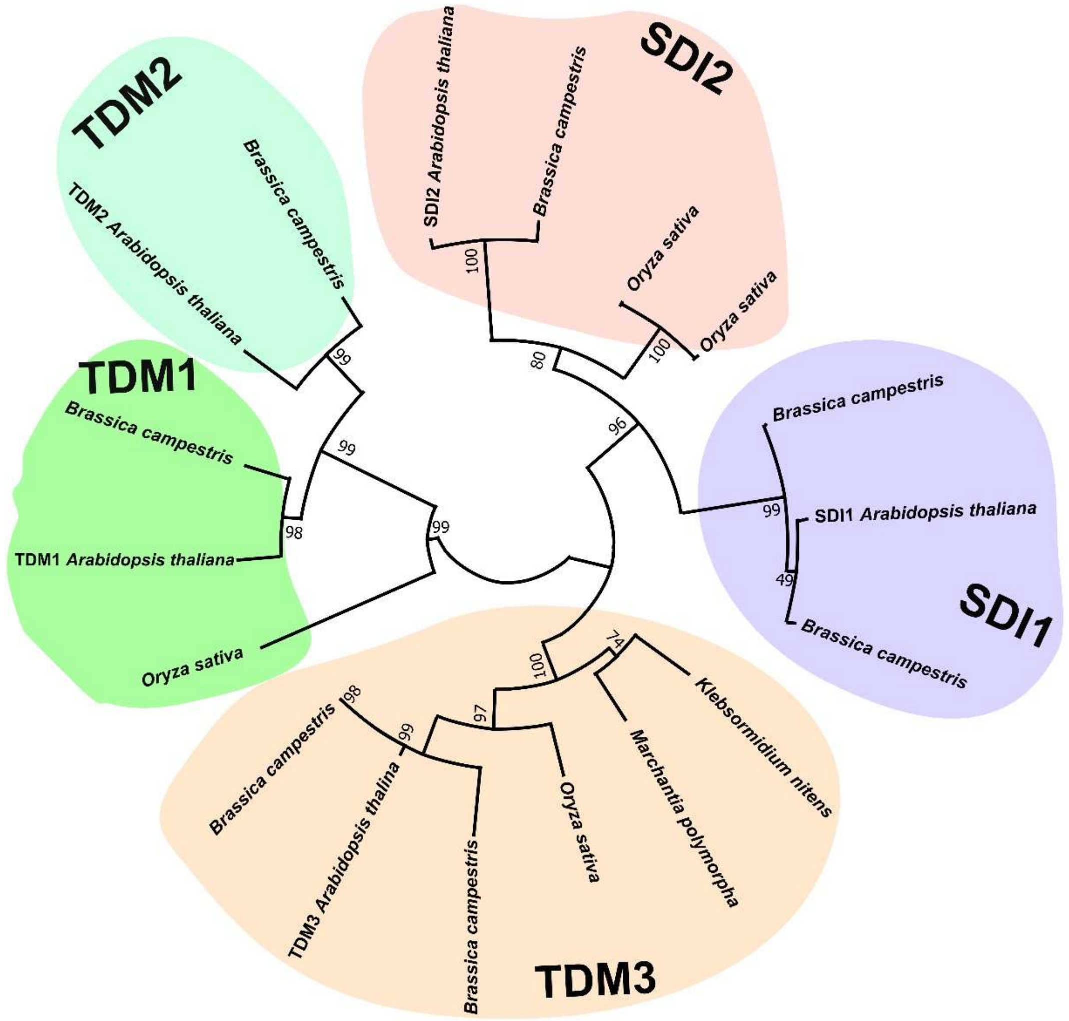


**Extended Data Fig. 2 | Expanded phylogenetic analysis of TDM/SDI proteins.** Phylogenetic analysis of TDM/SDI homologs from representative plant species illustrating the distribution of TDM1, TDM2, TDM3, SDI1 and SDI2 clades. The analysis includes homologs from *Arabidopsis thaliana*, *Brassica campestris*, *Oryza sativa*, *Marchantia polymorpha* and *Klebsormidium nitens*. Coloured areas indicate the major protein groups. Numbers at branch points indicate 1,000 bootstrap support values.

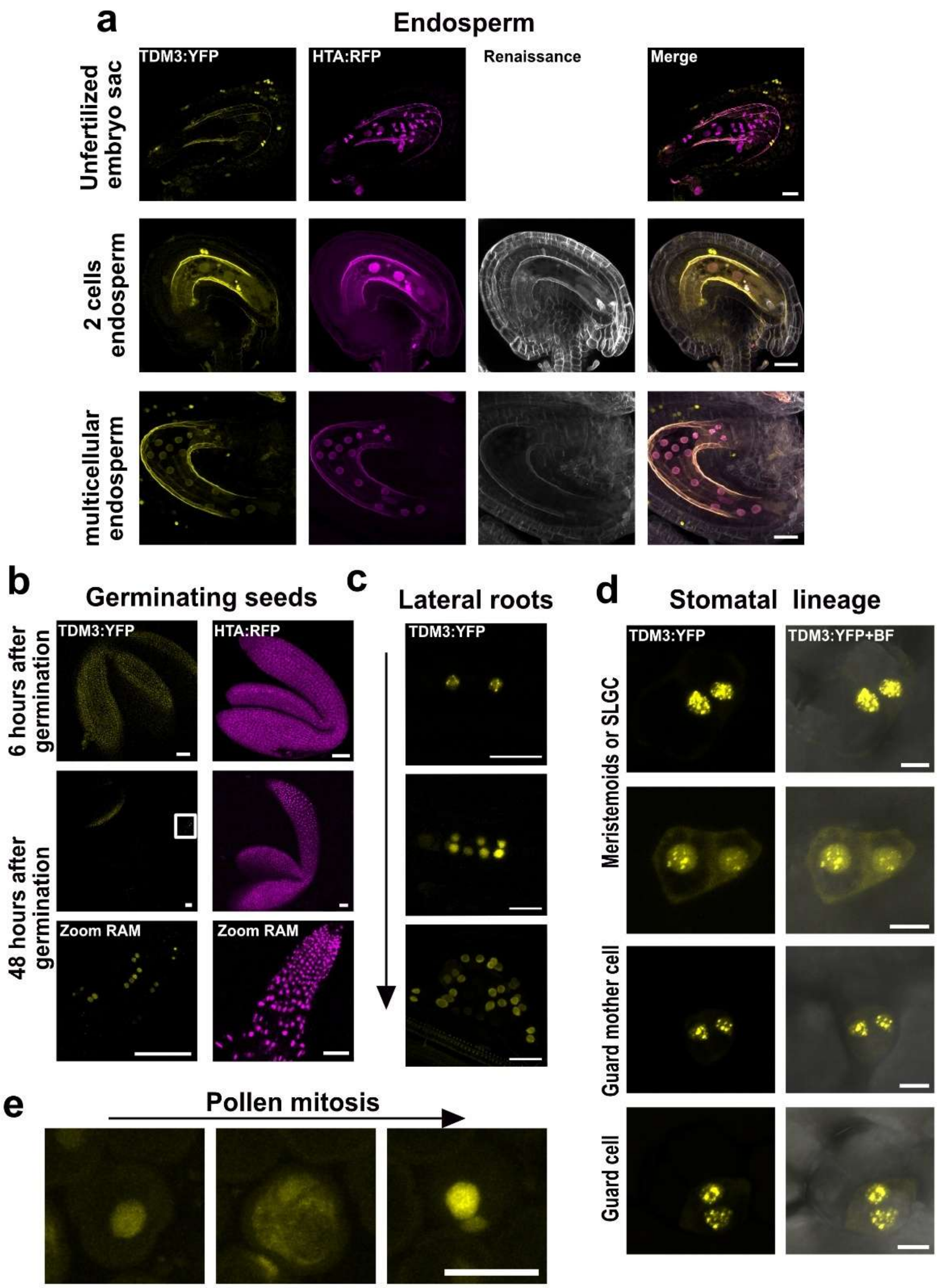


**Extended Data Fig.3 | TDM3 expression across different Arabidopsis tissues. a,** Representative MIP localization of TDM3:YFP (yellow) during endosperm development before pollination and at the two-cell and multicellular stages in ovules fertilized with pollen from plants expressing the histone marker HTA:RFP (violet). Renaissance staining is shown in grey. In addition to the developing endosperm, TDM3:YFP expression is also apparent in mitotically dividing cells of the maternal sporophitic integuments surrounding the developing zygotic tissue. **b,** Representative MIP images showing localization of TDM3:YFP during seed germination at 6 h and 48 h after imbibition together with HTA:RFP. No TDM3:YFP signal was detected at 6 h after imbibition. At 48 h, both TDM3:YFP and HTA:RFP signals were detected. HTA:RFP showed broad expression, whereas TDM3:YFP expression was restricted to a subset of cells in the root apical meristem. **c,** Representative MIPs showing expression of TDM3:YFP during lateral root development. The arrow indicates progression from early to late developmental stages. **d,** Representative MIPs showing localization of TDM3:YFP in stomatal lineage cells, including meristematic proliferative cells, guard mother cells and guard cells. Bright filed (BF) signal is shown where indicated. **e,** Representative MIP localization of TDM3:YFP during the first pollen division. Scale bars, 10 µm (**a**,**e**), 50 µm (**b**), 20 µm (**c**), and 5 µm (**d**).

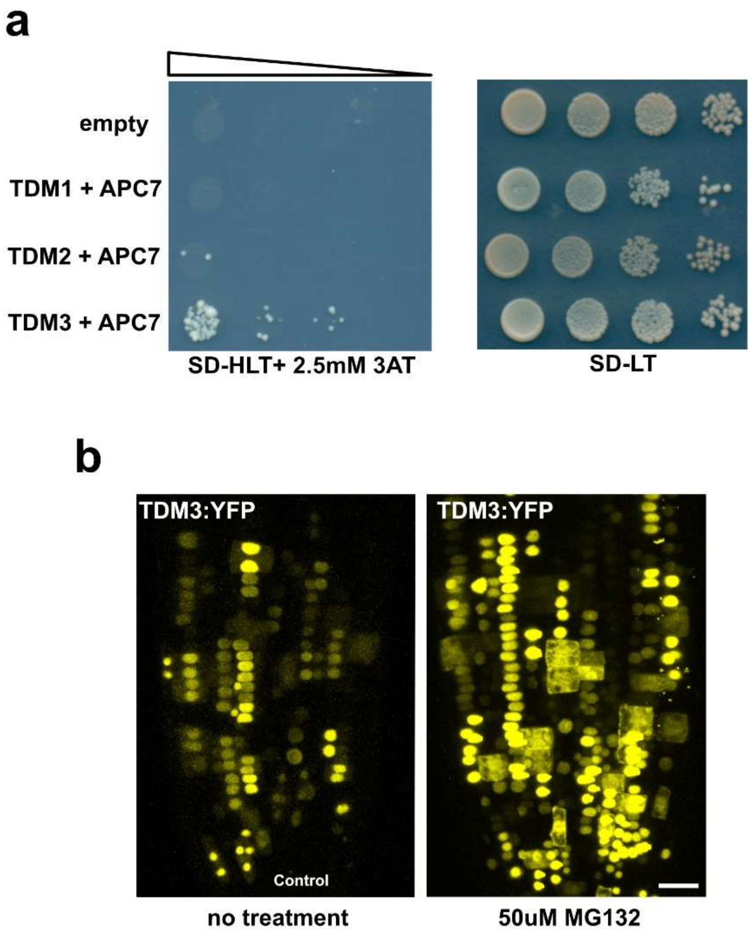


**Extended Data Fig.4 | TDM3 is subject to APC/C-dependent degradation. a,** Ten-fold dilution series of TDM1, TDM2 and TDM3 bait constructs analysed by yeast two-hybrid assay using APC7, CDC27 and CCS52 as prey constructs. Interactions were assessed on SD-HLT medium supplemented with 2.5 mM 3-AT and SD-LT was used as growth control. **b,** Accumulation of TDM3:YFP in the root apical meristem following treatment with the proteasome inhibitor MG132 (50 μM, 1 h) compared with untreated control. Scale bar, 20 μm (**b**).

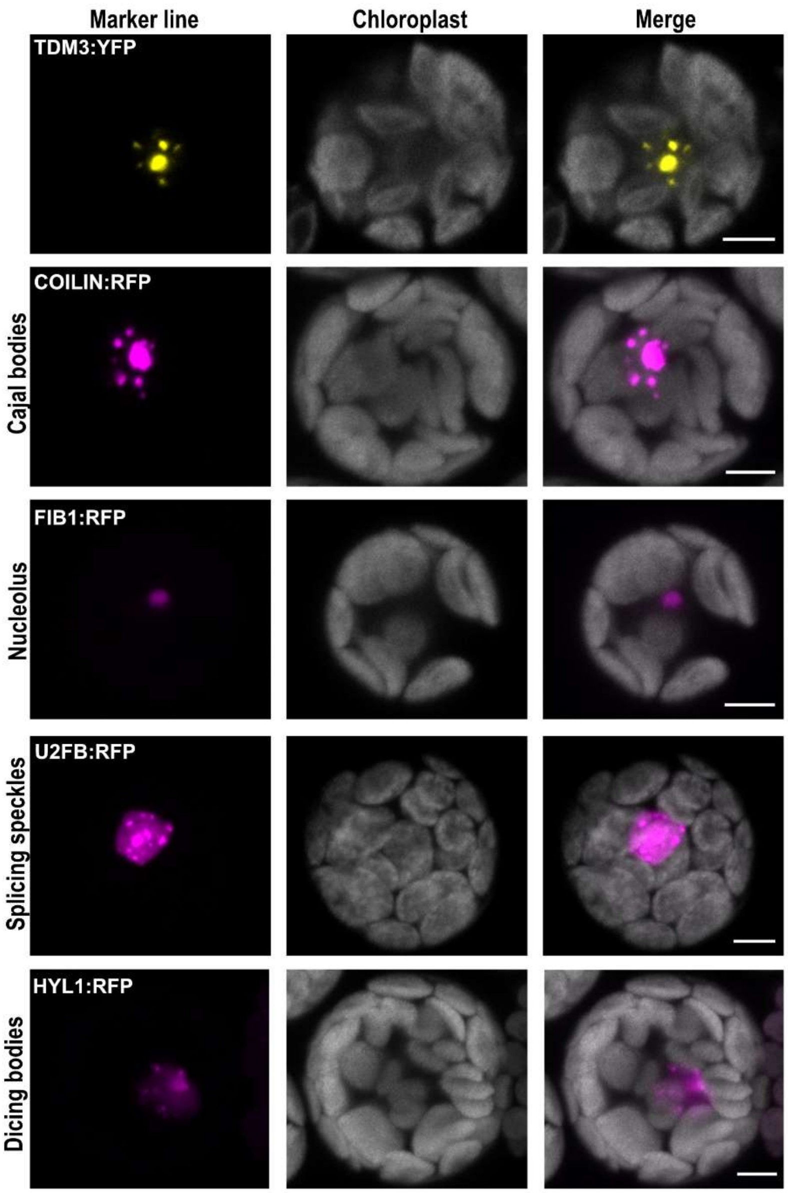


**Extended Data Fig. 5 | Expression of nuclear markers used for colocalization analysis.** Representative MIP of individual expression of TDM3 (*35S::TDM3:YFP*), COILIN (*35S::COILIN:RFP*), FIBRILARIN1 (*35S::FIB1:RFP*), U2AF35B (*35S::U2AF35B:RFP*), and HYL1 (*35S::HYL1:RFP*) in *Arabidopsis* mesophyll protoplasts. Scale bars, 5 µm.

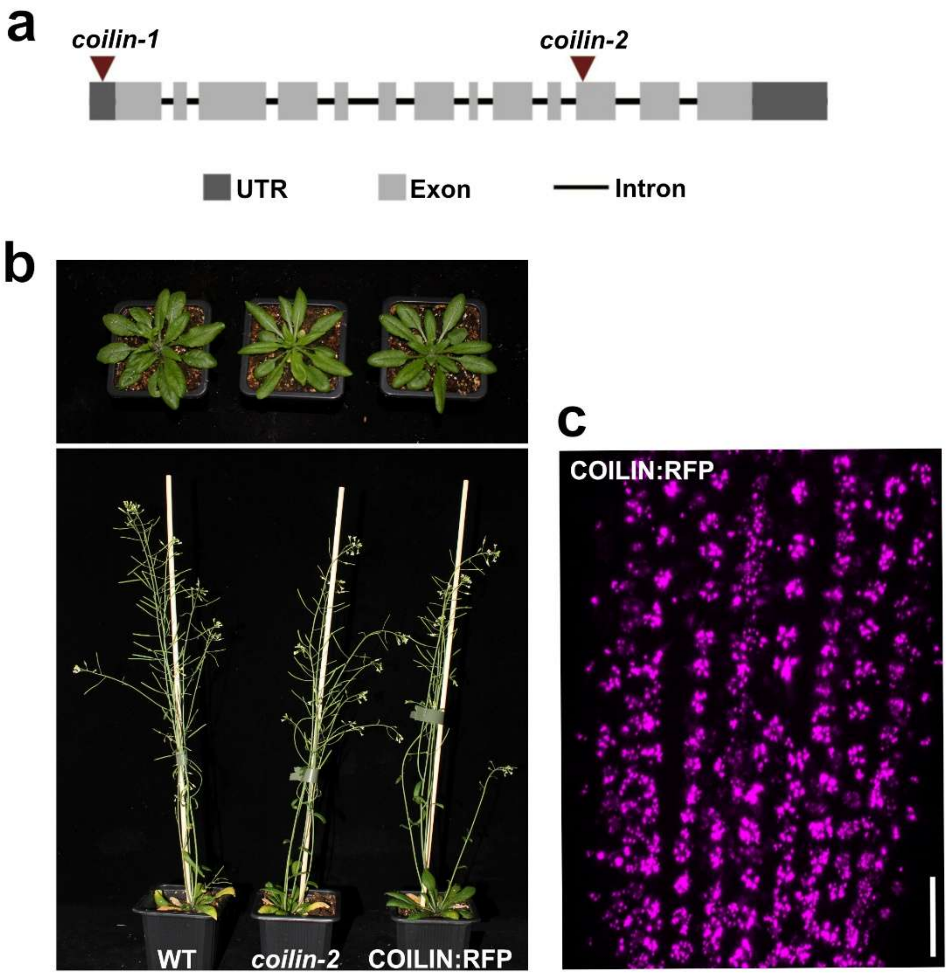


**Extended Data Fig. 6 | Characterization of COILIN. a,** Schematic representation of the *COILIN* gene with the indicated T-DNA insertion sites in *coilin-1* and *coilin-2* alleles. **b,** Comparison of leaf rosettes and 6 weeks old wild type plants, *coilin-2* mutants, and *coilin-2* mutants complemented with the *pCOILIN::COILIN:RFP* construct. **c,** Representative MIP showing localization of COILIN:RFP as a marker for Cajal bodies in root meristem. Scale bar, 20 µm (**c**).

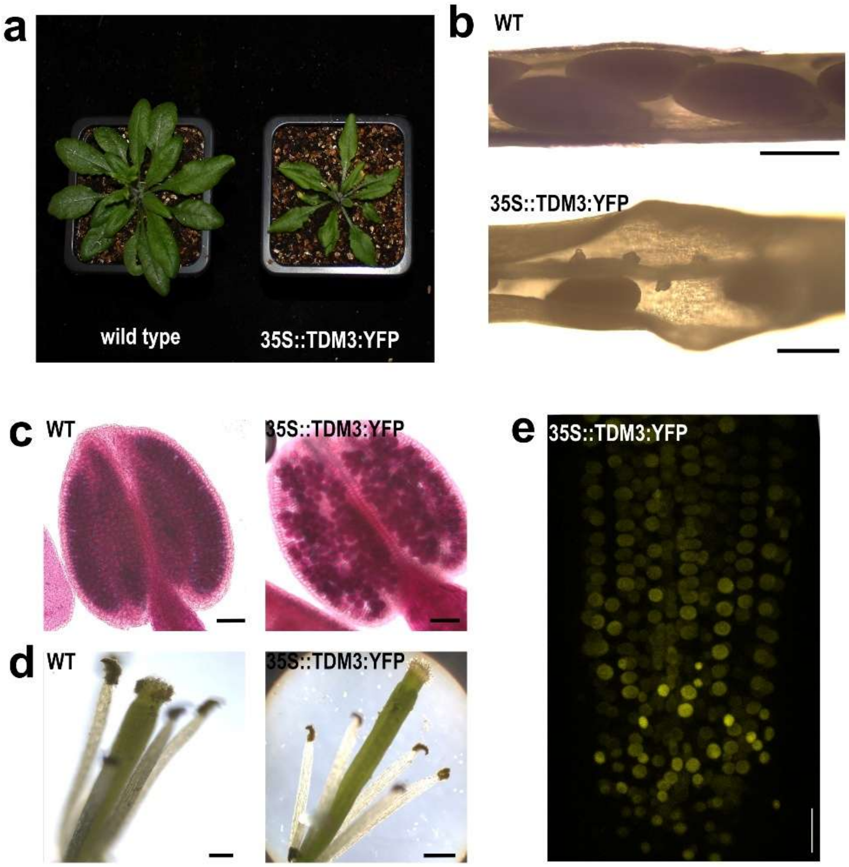


**Extended Data Fig. 7 | Characterization of plants harboring *35S::TDM3:YFP* construct overexpressing TDM3 protein**. Transgenic plants overexpressing TDM3 exhibited impaired growth as indicated by rosette morphology (**a**) and partial fertility. **b,** Microscopic view of developing siliques shows a number of aborted ovules. **c,** Alexander staining of anthers indicates reduced number of pollen. **d,** Stamens in mature flowers of *35S::TDM3:YFP* plants are shorter, which may lead to reduced pollination. **e,** Representative MIP showing localization of TDM3:YFP in the root apical meristem of *35S::TDM3:YFP* plant. Scale bar, 500 μm (**b**) 1mm (**c, d**), and 20 μm (**e**).

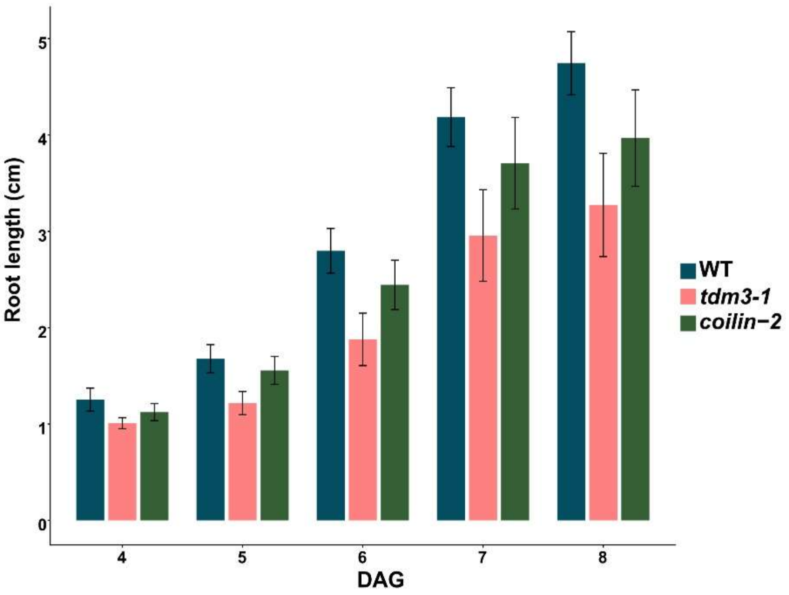


**Extended Data Fig. 8 | Root growth analysis of WT, *tdm3-1* and *coilin-2* seedlings.** Root length measurements are shown as mean ± s.d. at indicated developmental time points. Root growth trends were further supported by area under the curve (AUC) analysis, indicating reduced cumulative root growth in *tdm3-1* compared with WT (Cumulative AUC analysis: WT = 11.66; *tdm3-1* = 8.19, $P$ = 0.0354; *coilin-2* = 10.25, $P$ = 0.3529).

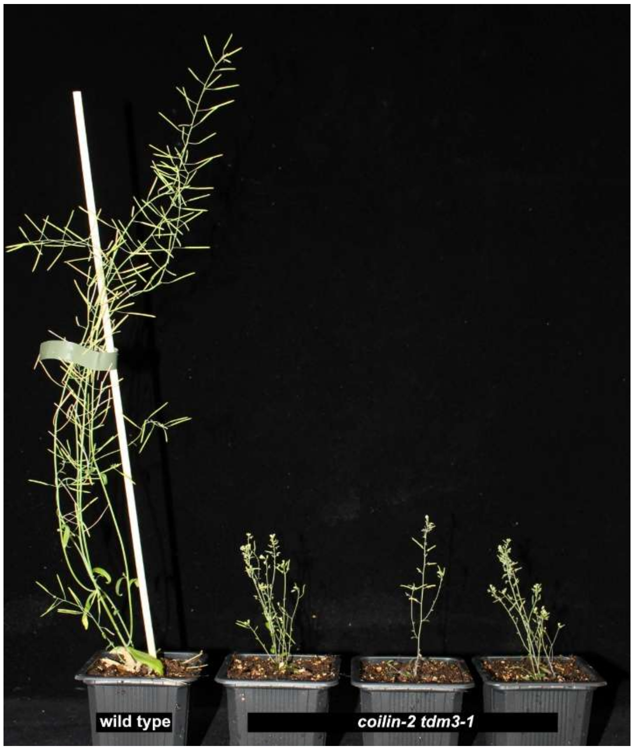


**Extended Data Fig. 9 | Genetic interaction between *tdm3-1* and *coilin-2*.** Phenotypic comparison of eight weeks old wild type and *tdm3-1 coilin-2* double mutants.

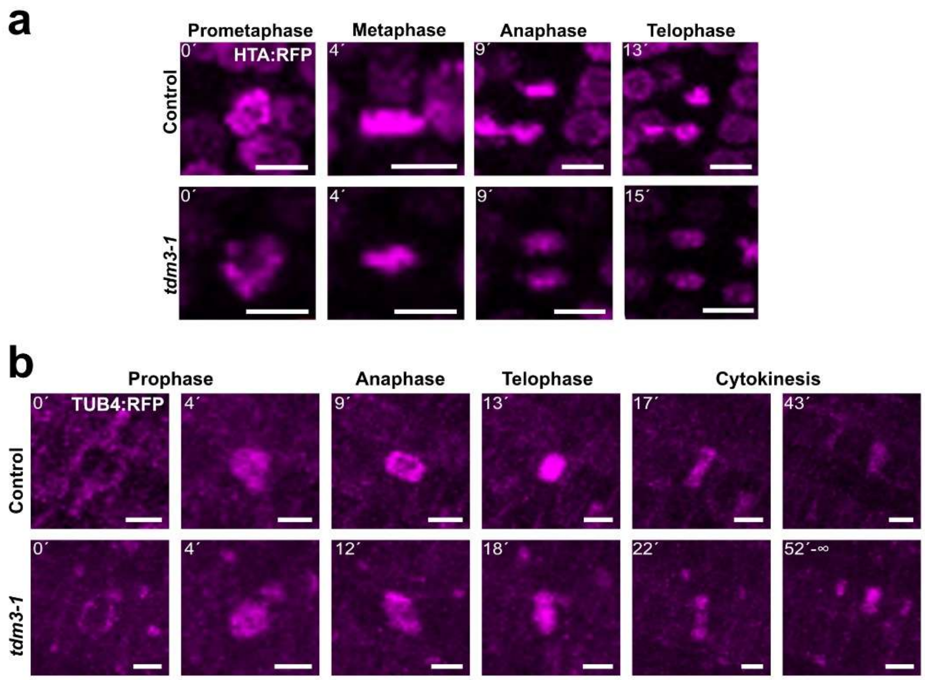


**Extended Data Fig. 10 | Analysis of mitotic progression using HTA:RFP and TUB4:RFP markers.** Representative time-lapse images of root cells undergoing mitosis and expressing **a,** HTA:RFP (histone marker) or **b,** TUB4:RFP (tubulin marker). Images show the mitotic stages used for the quantification of mitotic progression. Scale bars, 5 µm. Numbers indicate the elapsed time in minutes relative to nuclear envelope breakdown.

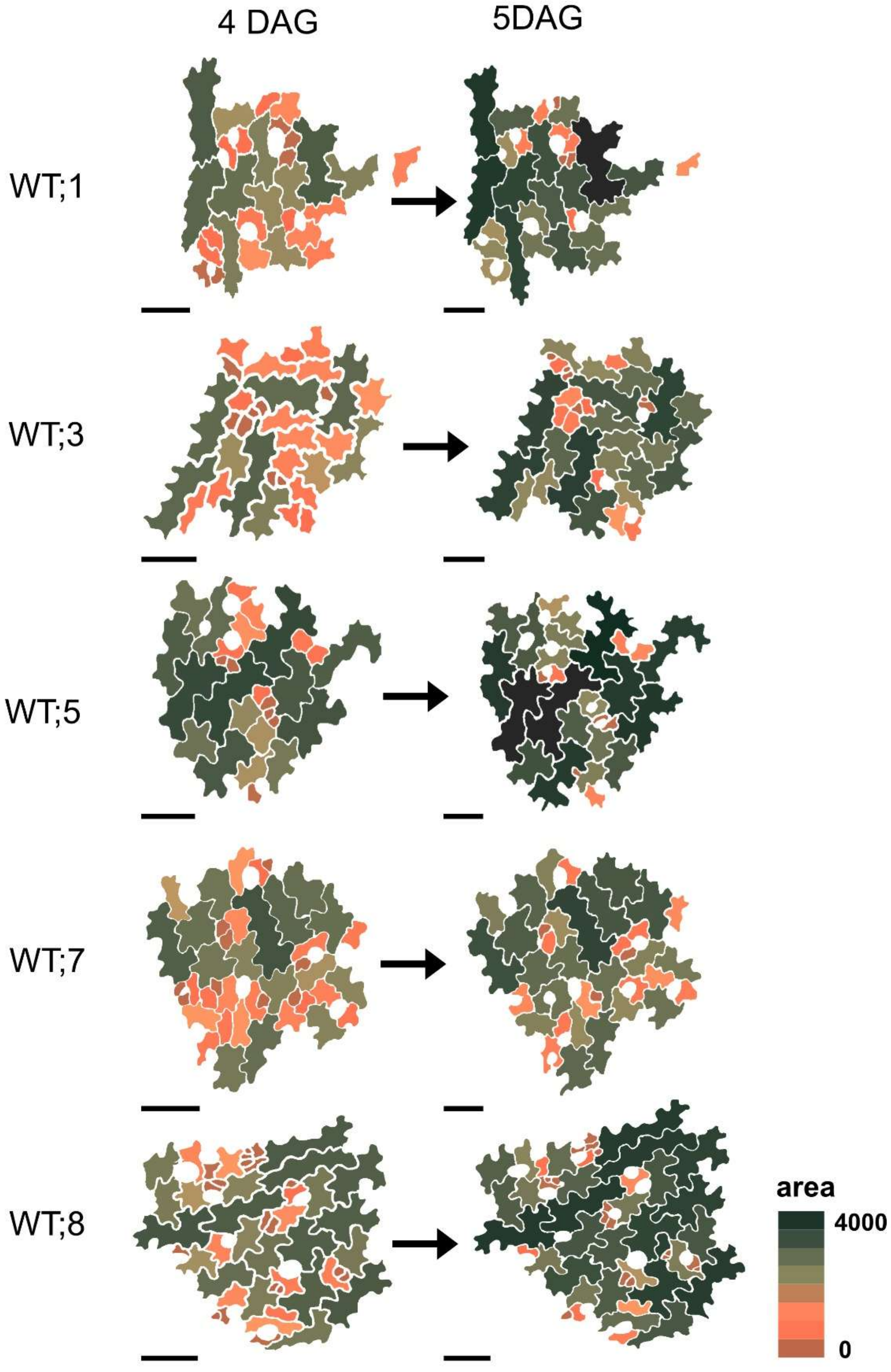


**Extended Data Fig. 11 | Spatial maps depicting the epidermal cells in wild type cotyledons in the same area at 4 and 5 DAG.** These datasets were used for quantifications presented in Fig. 3f–g. The heat map indicates cell area (μm²). Scale bar= 50 μm.

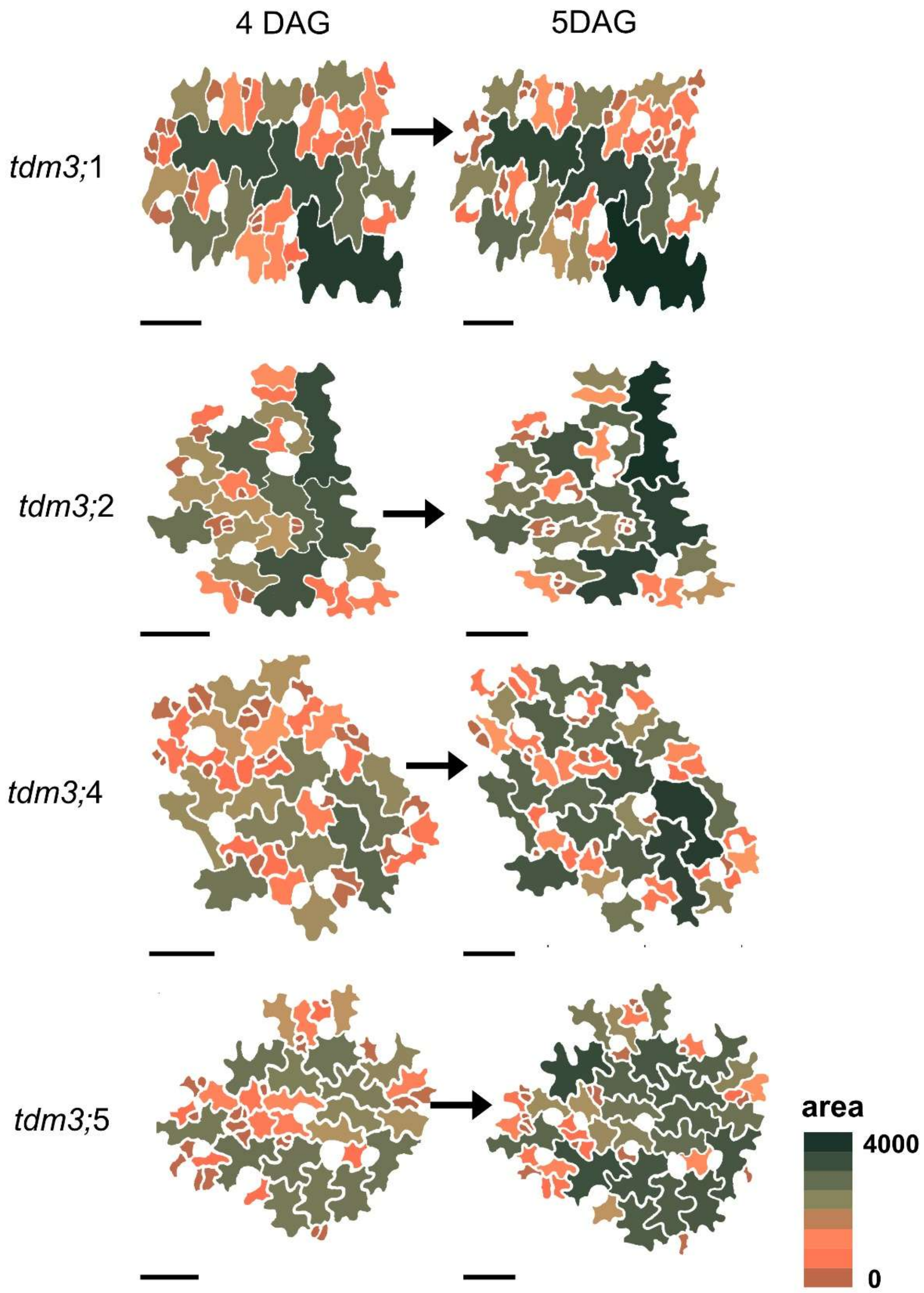


**Extended Data Fig. 12 | Spatial maps of epidermal cells in *tdm3-1* cotyledons imaged in the same area at 4 and 5 DAG.** These datasets were used for the quantifications presented in Fig. 3f,g. The heat map indicates cell area (μm²). Scale bar= 50 uM.

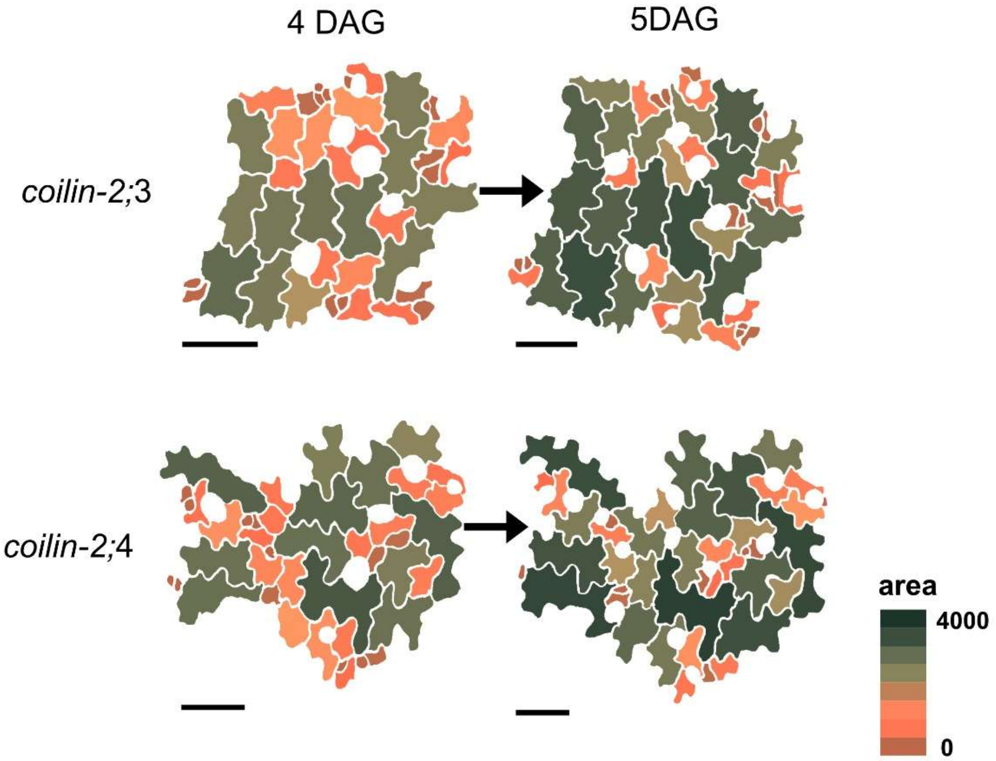


**Extended Data Fig. 13 | Spatial maps of epidermal cells in *colin-2* cotyledons imaged in the same area at 4 and 5 DAG.** These datasets were used for the quantifications presented in Fig. 3f,g. The heat map indicates cell area (μm²). Scale bar= 50 uM.

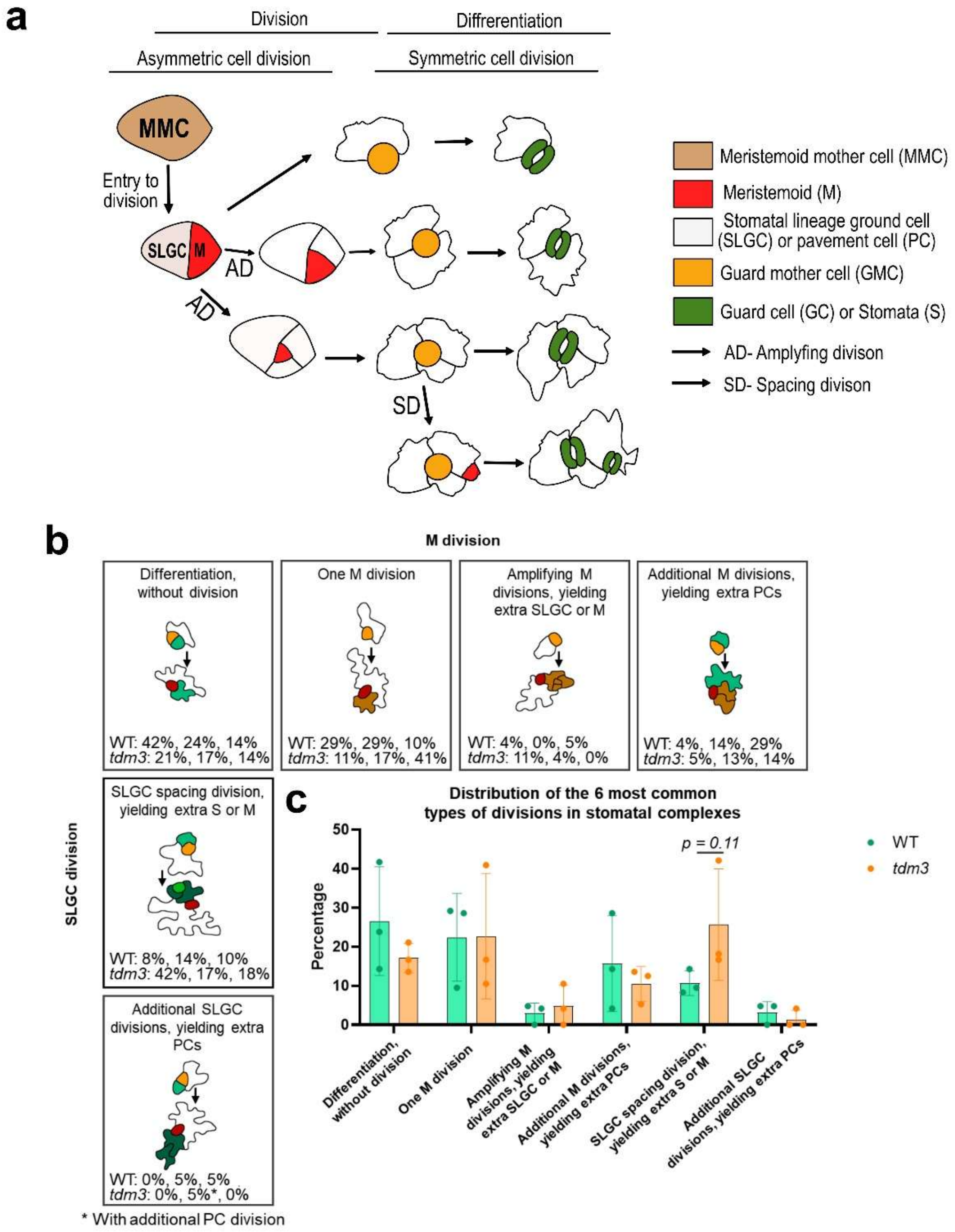


**Extended Data Fig. 14 | Analysis of stomatal progression in true leaves of *tdm3-1*. a,** Schematic representation of stomatal and pavement cell lineage development, showing asymmetric entry division that generates meristemoids (M) and stomatal lineage ground cells (SLGCs), followed by amplifying and spacing divisions that lead to the differentiation of guard and pavement cells. **b,** Representative stomatal division scenarios identified during lineage tracking of epidermal cells in developing true leaves between 10 and 21 DAG. The six most frequent division types within stomatal complexes lineages are shown. **c,** Percentage distribution of the six division types in wild type and *tdm3* leaves.

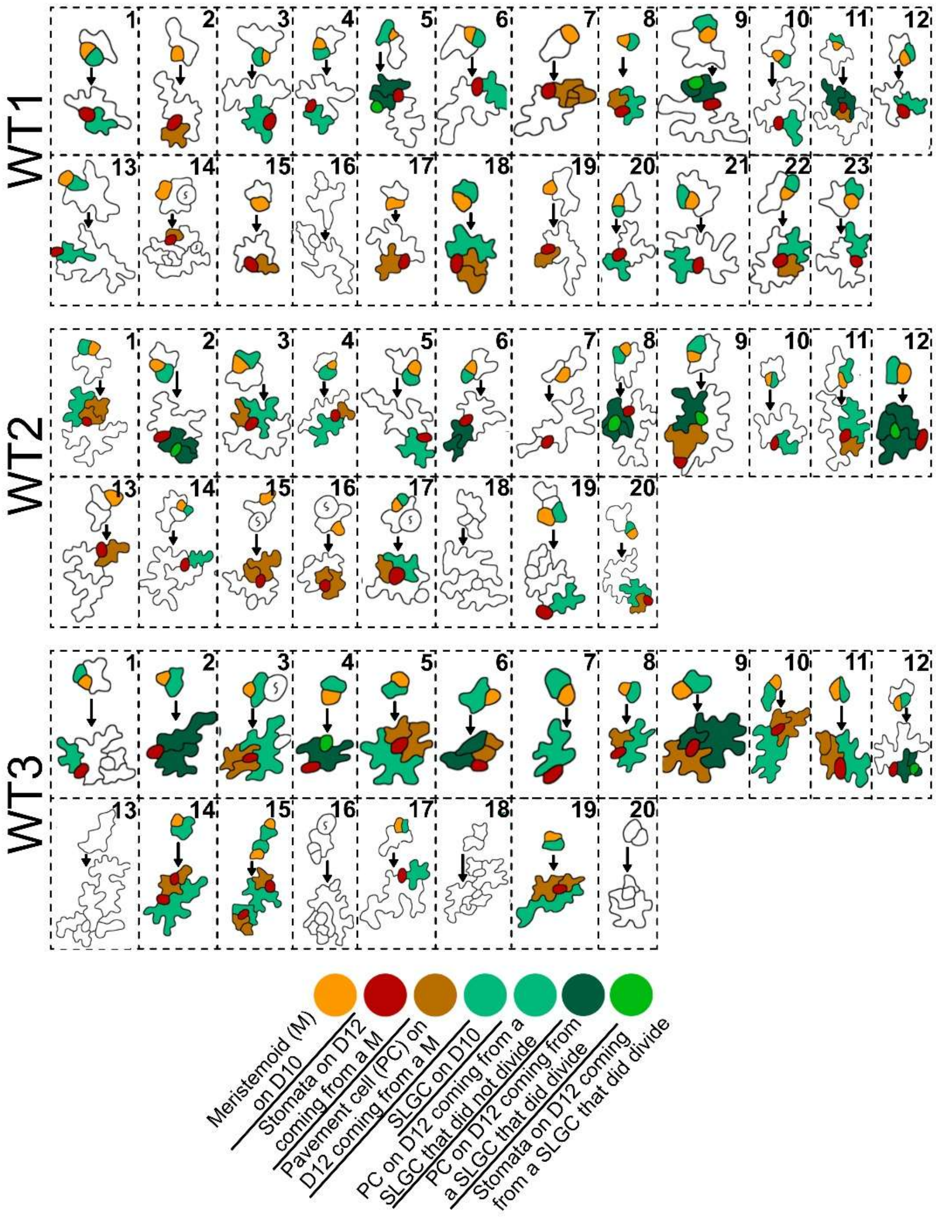


**Extended Data Fig. 15 | Stomatal complexes used for tracking division types in wild type true leaves.** Each box shows a stomatal complex at 10 DAG and the cells derived from this complex at 21 DAG. The analysis was performed on three independent biological samples (WT1, WT2, WT3). Colors indicate distinct stomatal lineage cell identities, including meristemoids (M), stomatal lineage ground cells (SLGCs), pavement cells (PCs), and guard cells (GCs).

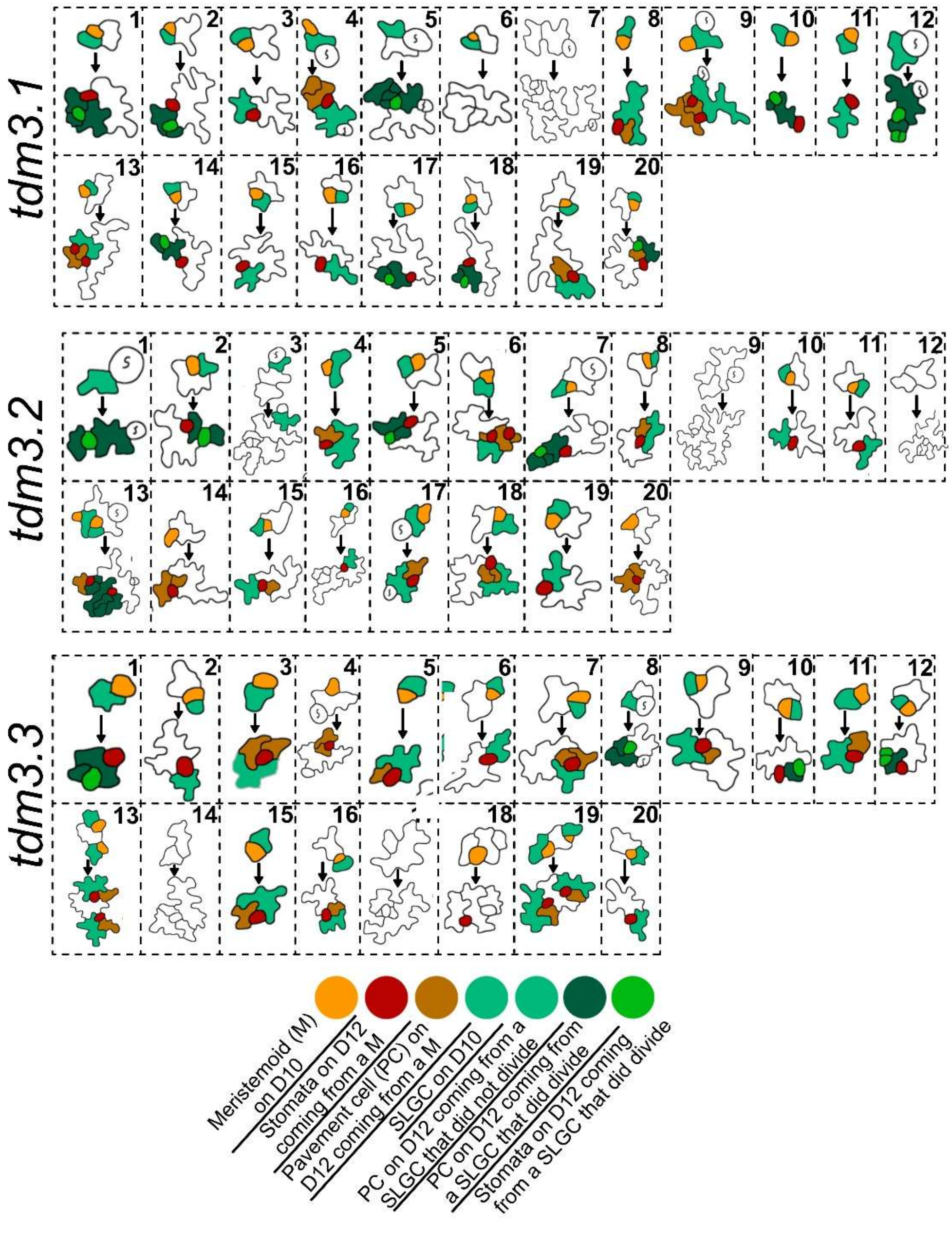


**Extended Data Fig. 16 | Stomatal complexes used for tracking division types in *tdm3-1* true leaves.** Each box shows a stomatal complex at 10 DAG and the cells derived from this complex at 21 DAG. The analysis was performed on three independent biological samples (*tdm3.1*, *tdm3.2, tdm3.3*). Colors indicate distinct stomatal lineage cell identities, including meristemoids (M), stomatal lineage ground cells (SLGCs), pavement cells (PCs), and guard cells (GCs).

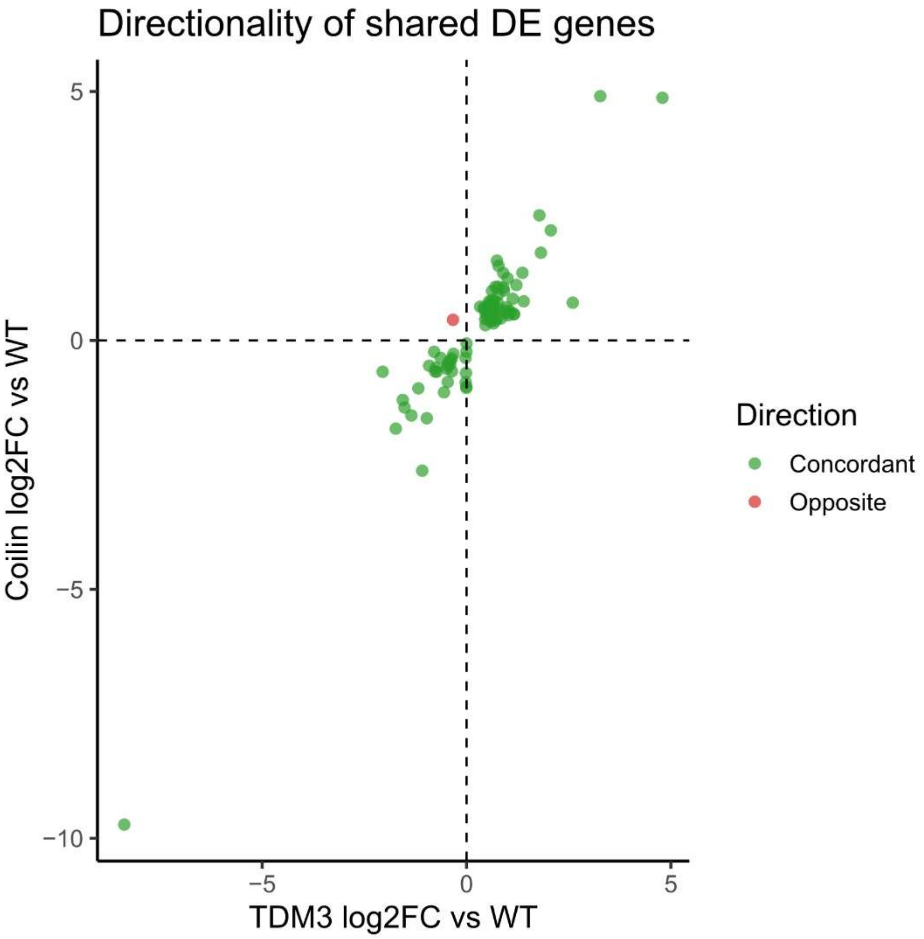


**Extended Data Fig. 17 | Shared transcriptional responses in *tdm3-1* and *coilin-2*.** Scatter plot showing the correlation of shared differentially expressed genes (DEGs) between *tdm3-1* and *coilin-2*. Log2 fold change values for *tdm3-1* and *coilin-2* are shown on the x and y axes, respectively. Each point represents a single gene. The strong linear relationship indicates concordant transcriptional responses between the two genotypes.

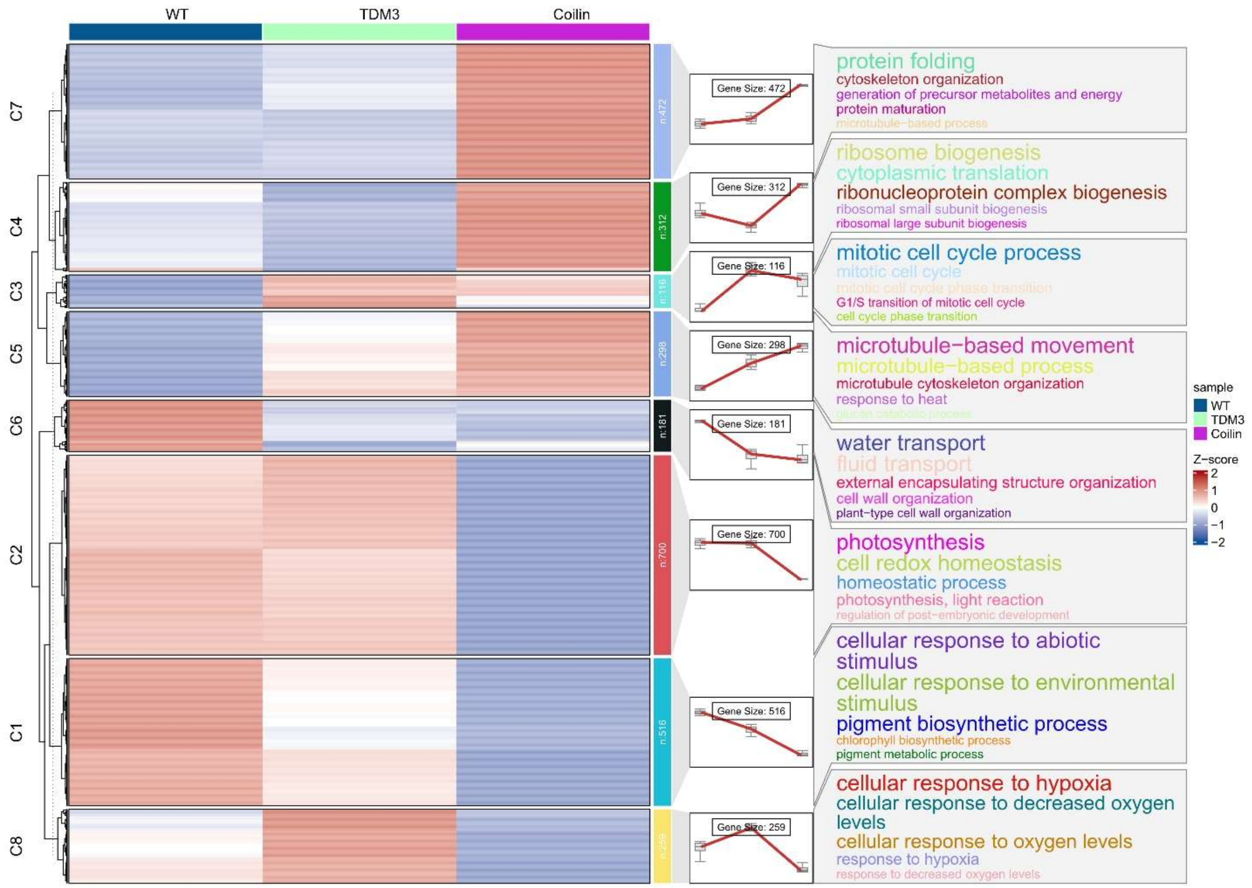


**Extended Data Fig. 18 | Co-expression clustering and cell-cycle dynamics.** Heat map showing k-means clustering (k = 8) of a set of 2,855 differentially expressed genes in *tdm3-1* and *coilin-2*. Data represent Z-score–normalized, replicate-averaged variance-stabilizing transformed (VST) counts. The number of genes in each cluster and the most significantly enriched gene ontology (GO) terms for each cluster are indicated.

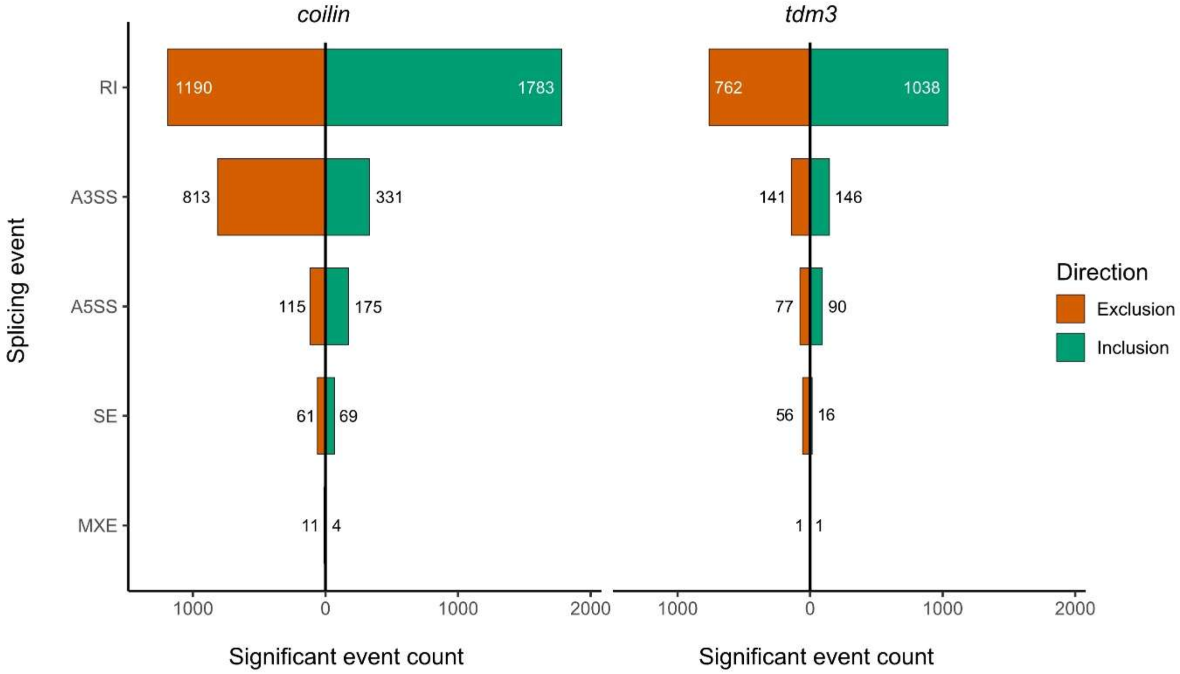


**Extended Data Fig. 19 |** Directionality of splicing events. Significant events are partitioned into Inclusion (retention of alternative sequence) or Exclusion (skipping of alternative sequence). Note the strong bias toward inclusion in the RI category for both genotypes.

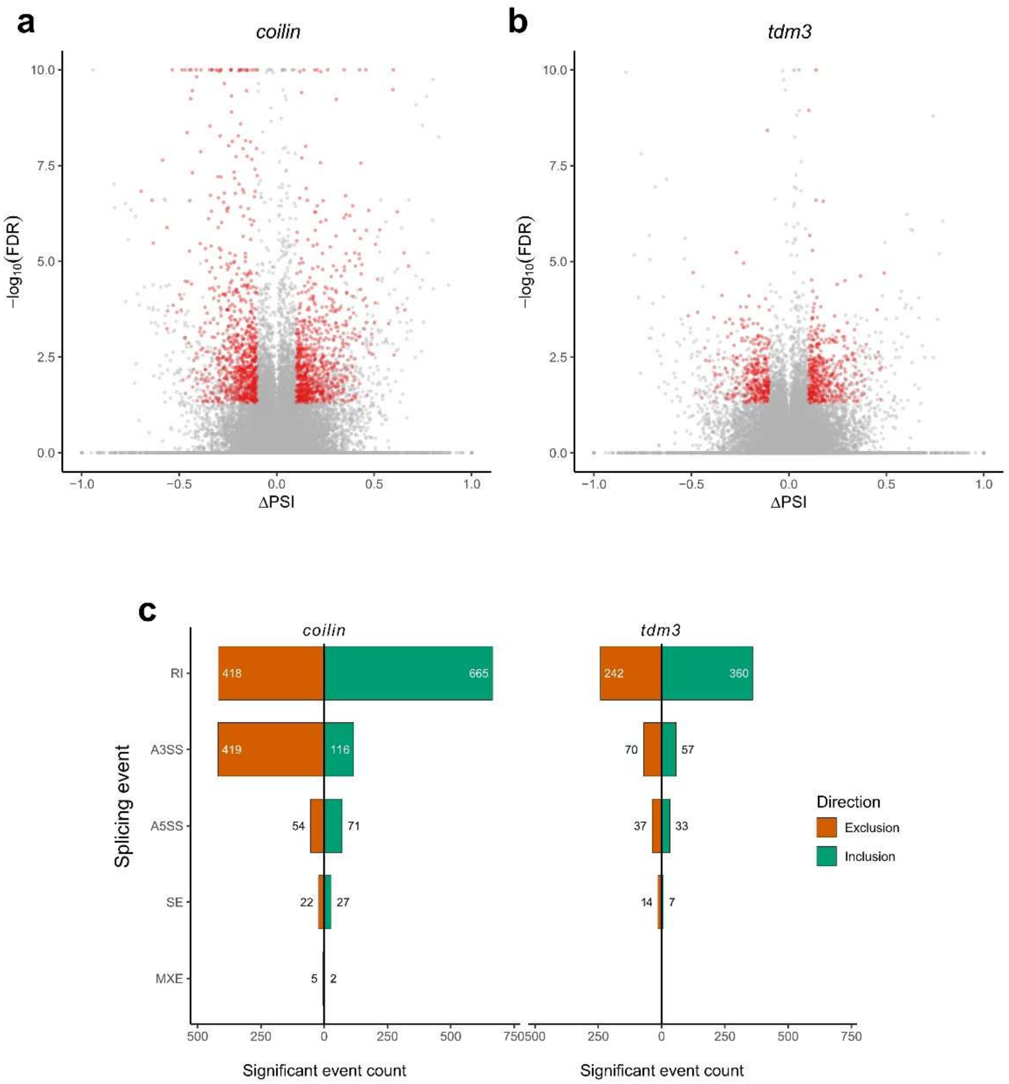


**Extended Data Fig. 20 | Alternative splicing changes in *coilin-2* and *tdm3-1*. a,b,** Volcano plot of differential alternative splicing events in *coilin-2* (**a**) and *tdm3-1* (**b**), showing ΔPSI (Percent Spliced In) versus −log10(FDR). Significant events (FDR < 0.05, |ΔPSI| > 0.1, read count > 10) are highlighted in red.**c,** Diverging bar plots showing the numbers of inclusion (ΔPSI > 0.1; green) and exclusion (ΔPSI < −0.1; orange) events across alternative splicing categories.
